# Characterization of *Poldip2* knockout mice: avoiding incorrect gene targeting

**DOI:** 10.1101/2021.02.02.429447

**Authors:** Bernard Lassègue, Sandeep Kumar, Rohan Mandavilli, Keke Wang, Michelle Tsai, Dong-Won Kang, Marina S. Hernandes, Alejandra San Martín, Hanjoong Jo, W. Robert Taylor, Kathy K. Griendling

## Abstract

POLDIP2 is a multifunctional protein whose roles are only partially understood. Our laboratory previously reported physiological studies performed using a mouse gene trap model, which suffered from two limitations: perinatal lethality in homozygotes and constitutive *Poldip2* inactivation. To overcome these limitations, we developed a new conditional floxed *Poldip2* model. The first part of the present study shows that our initial floxed mice were affected by an unexpected mutation, which was not readily detected by Southern blotting and traditional PCR. It consisted of a 305 kb duplication around *Poldip2* with retention of the wild type allele and could be traced back to the original targeted ES cell clone. We offer simple suggestions to rapidly detect similar accidents, which may affect genome editing using both traditional and CRISPR-based methods. In the second part of the present study, correctly targeted floxed *Poldip2* mice were generated and used to produce a new constitutive knockout line by crossing with a Cre deleter. In contrast to the gene trap model, many homozygous knockout mice were viable, in spite of having no POLDIP2 expression. To further characterize the effects of *Poldip2* ablation in the vasculature, an RNA-seq experiment was performed in constitutive knockout carotid arteries. Results support the involvement of POLDIP2 in multiple cellular processes and provide new opportunities for future in-depth study of its functions.

## Introduction

POLDIP2 is a protein with a surprisingly large variety of roles, affecting apparently unrelated cellular processes and physiological functions. This pleiotropism of POLDIP2 results in part from its expression in most tissues [1–12] and subcellular compartments, including the nucleus, cytosol and mitochondria [13–15]. In addition, POLDIP2 can potentially bind to numerous other proteins, currently nearly 200, according to its NCBI human gene record. Since most of the interactions listed in the NCBI database have not yet been characterized, additional roles of POLDIP2 will likely be discovered in the future. Below is a brief summary of POLDIP2 functions known to date. More detail can be found in a recent review by Hernandes et al. [16].

Initial studies, carried out in vitro, revealed that POLDIP2 interacts with the p50 subunit of DNA polymerase δ [17], hence its name of POLymerase Delta Interacting Protein 2. However, further work showed that, in addition to polymerase δ, POLDIP2 also activates other polymerases involved in DNA repair (translesion synthesis) [18–22]. This role of POLDIP2 in the DNA damage response has just been reviewed in detail [23]. In a related set of functions, POLDIP2 favors cell division and proliferation through its effects on the mitotic spindle [24], as well as cell cycle activators and inhibitors [25, 26]. These results may explain the association of POLDIP2 with various types of cancer [27–29].

Additional cellular studies, using overexpression and knockdown, brought to light striking effects of POLDIP2 on the cytoskeleton mediated by Ect2 and RhoA [10, 30]. POLDIP2 increases stress fiber formation and stabilizes focal adhesions [10, 31]. It also intensifies the interaction of actin and vinculin [32], and mediates TGFβ-induced smooth muscle alpha actin expression [11]. These regulations can have important consequences on cell migration [10, 30–32]. Additionally, an inhibitory effect of POLDIP2 on autophagy and the proteasome has been implicated in tau aggregation in a model Alzheimer’s disease [8].

The observation that POLDIP2 can be highly expressed in mitochondria [15, 33, 34] led to the discovery of its positive effect on the Krebs cycle, through inhibition of the mitochondrial ClpXP protease [35]. Due to its impact on mitochondria and nuclear transcription factors, POLDIP2 can regulate differentiation and metabolism [5, 36].

Many of the effects of POLDIP2 appear to be mediated by reactive oxygen species, especially H2O2 generated by the NADPH oxidase Nox4. POLDIP2 can associate with the p22phox subunit [9, 10] and activate the oxidase [10] or induce Nox4 upregulation [11]. This positive effect of POLDIP2 on H2O2 has been observed not only in cells [5, 10, 11, 30–32], but also in vivo, using the mouse gene trap model introduced by our laboratory [12, 37–39].

This mouse gene trap model allowed in vivo studies of physiopathological functions of POLDIP2 to be carried out for the first time. Thus, it was found that POLDIP2 promotes neovascularization in a model of hindlimb ischemia via ROS-dependent activation of metalloproteinases [37]. POLDIP2 also preserves vascular structure and function by preventing excessive accumulation of extracellular matrix, including collagen and fibronectin, thereby enhancing vessel contraction [12]. The mechanism of this extracellular matrix reduction by POLDIP2, studied in detail in cultured vascular smooth cells [40], may explain the antifibrotic activity of the plant alkaloid matrine [41]. However, an opposite finding in renal fibroblasts suggests that POLDIP2 can mediate TGFβ-induced fibrosis [11].

Additional adverse effects of POLDIP2 in disease have been uncovered using the gene trap model. Thus, *Poldip2^+/-^* mice are protected against blood brain barrier leakage and subsequent harmful edema and inflammation after an experimental ischemic stroke [4] or LPS administration [7]. The integrity of the blood brain barrier was also preserved by intracerebral administration of *Poldip2* siRNA in a mouse meningitis model [42]. Similarly, *Poldip2^+/-^* mice are protected against lung edema and inflammation after LPS [1]. These studies suggest that POLDIP2 is a promising therapeutic target for pathologies involving breakdown of a barrier, such as stroke and sepsis, for which new treatments are urgently needed. POLDIP2 is especially interesting because of its upstream position in signaling pathways leading to reactive oxygen species production, inflammation, cell migration and proliferation.

Further exploration of the multiple functions of POLDIP2 will require the development of new tools applicable to cells and organisms. Initial in vivo studies relied on the mouse gene trap model mentioned above, which suffered from two main limitations. First, *Poldip2* knockdown was constitutive, thereby affecting all cell types throughout the life of the animals [12]. All other *Poldip2* gene trap alleles currently listed in the Mouse Genome Informatics database (http://www.informatics.jax.org/allele/summary?markerId=MGI:1915061) are similarly constitutive. Second, because most homozygous gene trap mice died shortly after birth [12, 25], experiments were limited to heterozygous animals. Therefore, we sought to develop a conditional knockout model to allow finer spatial and temporal control of *Poldip2* expression in vivo, using the well-established Cre-lox system.

In the present study we targeted the mouse *Poldip2* gene, using a classical homologous recombination strategy in embryonic stem (ES) cells, with the help of a reputable commercial company. In spite of using the most accepted methods and quality control steps, our first line of mice was flawed. Characterization of its genetic defect revealed that a mishap occurred during gene targeting, with partial recombination of the targeting construct, duplication of adjacent genes and retention of the wild type allele. We believe this is an underreported pitfall and hope our experience will incite others to incorporate DNA copy number assays in future gene targeting projects, including those based on CRISPR-mediated genome editing.

After identifying a different clone of correctly targeted ES cells, we were able to produce the desired floxed *Poldip2* mice. To verify that the floxed allele can be inactivated by excision with Cre recombinase, we crossed floxed mice with a transgenic Cre deleter expressed in early embryos to produce a new line with a ubiquitous deletion. POLDIP2 expression was abolished in mice homozygous for the constitutively excised allele, showing that it is null, as desired. Interestingly, a much higher proportion of live homozygous knockout *Poldip2* mice could be obtained from this line than from the gene trap. We were thus able to perform initial characterization of their phenotype.

In summary, understanding the nature of the genetic defect that impeded our first gene targeting attempt allowed us to successfully produce two new lines of mice: a floxed conditional and a constitutive knockout, that will allow refined in vivo physiopathological studies of POLDIP2.

## Materials and Methods

### Genetically modified mice

Vector construction and traditional gene targeting were performed by genOway (Lyon, France). The targeting construct included positive (neomycin) and negative (diphtheria toxin A) selection cassettes. Segments of the *Poldip2* gene were amplified using 15-20 cycles of PCR with Accuprime Taq DNA polymerase HF (Invitrogen) from C57BL/6 genomic DNA. Homology regions included a long 4.5 kb 5’ arm encompassing exons 2 and 3, and a short 3 kb 3’ arm spanning exons 5-9. Exon 4 and the neomycin cassette (itself flanked by FRT sites) were surrounded by loxP sites. Recombination of loxP sites by Cre was designed to delete 1.2 kb of genomic DNA, including exon 4 with its 97 bp of coding region, thereby producing a frame shift and a premature stop codon in exon 5. The construct was linearized with Pme I, purified and transfected by electroporation into embryonic stem (ES) cells from C57BL/6 black mice. After 48 hours in culture, cells were incubated with 200 μg/ml G418 (positive selection), leading to the isolation of 257 antibiotic-resistant clones. Screening by PCR across the 3’ arm, using primers p1 and p2 (Table 1), identified 10 positive clones, 7 of which were confirmed by southern blotting using 5’ and 3’ external probes. Six positive clones were injected into C57BL/6J albino blastocysts and implanted into OF1 females, which produced 15 male chimeras (>50% black), including two from clone B11-G2 and four from clone B12-H5. Three male chimeras were crossed with C57BL/6 Flp deleter females to excise the neomycin cassette. Two heterozygous pups, both derived from ES cell clone B12-H5, were identified by PCR genotyping, verified by Southern blotting with the 3’ external probe and used as colony founders.

**Table 1.**
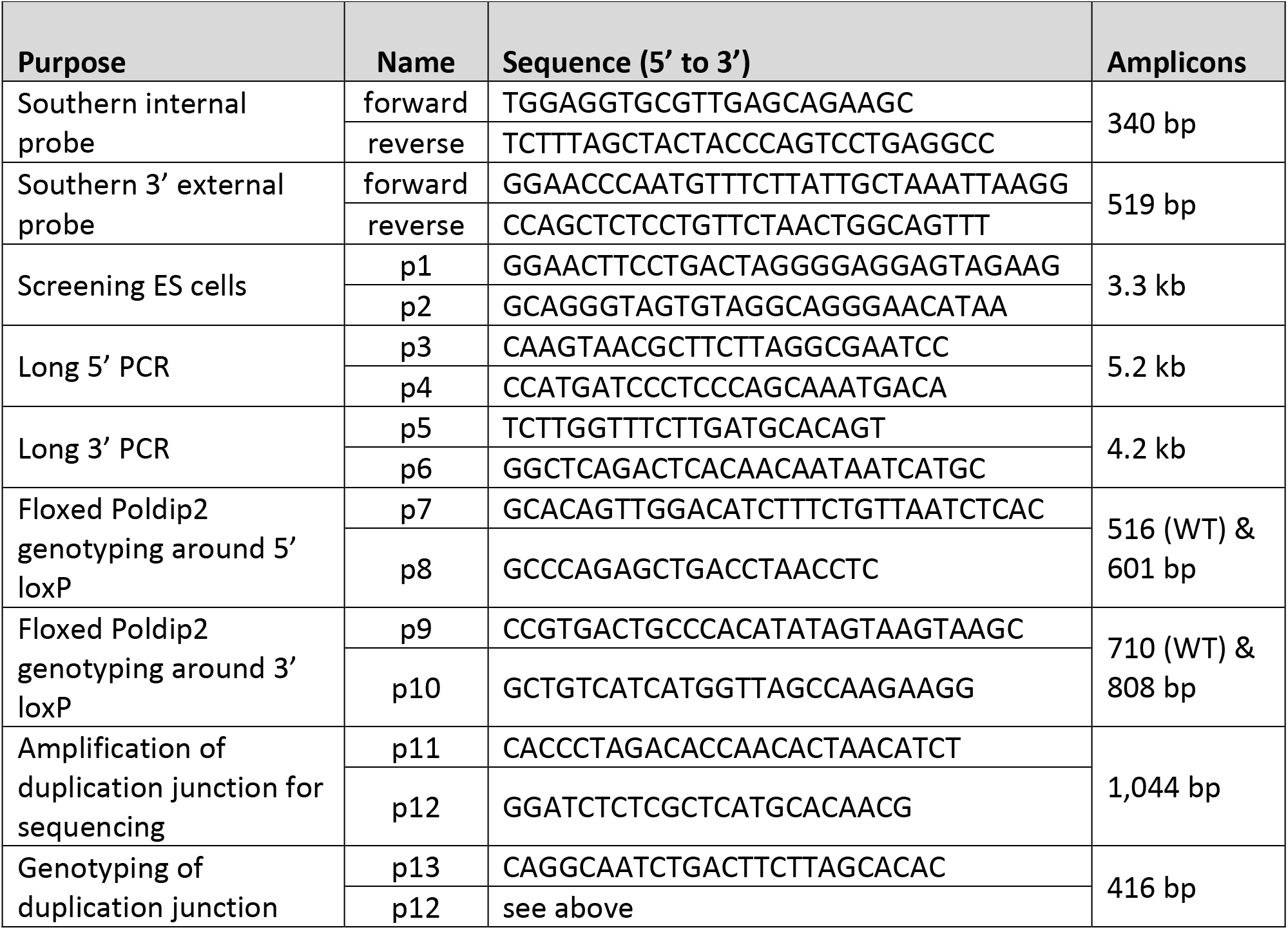
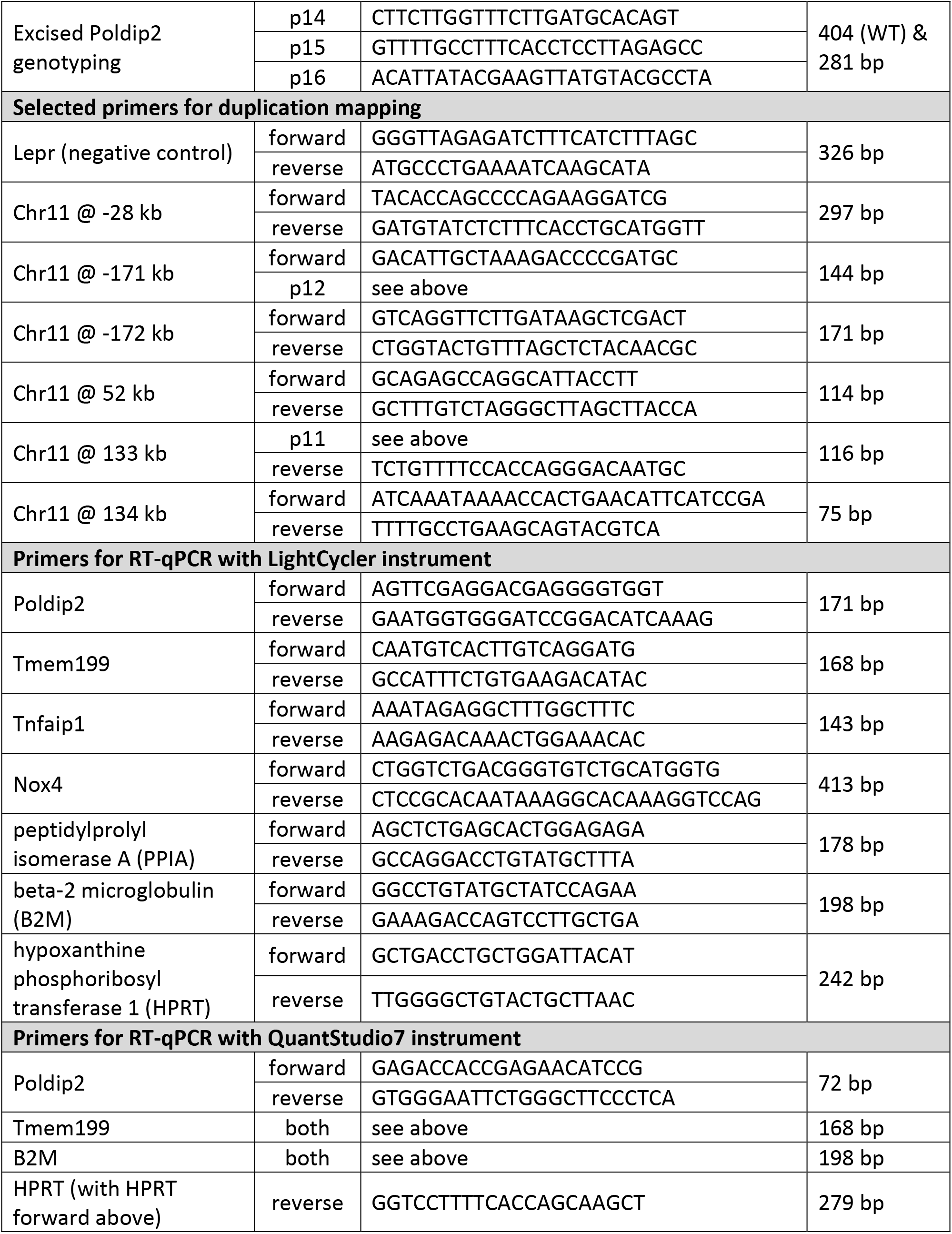
**PCR primers**

### Southern blots

Southern blotting protocols were designed and carried out by genOway, using DNA probes labeled with ^32^P. Probe sequences were assessed using BLAST software to ensure specificity in the murine genome and tested using wild-type genomic DNA. The following combinations of restriction enzymes and probes were used. Digestion with EcoRV followed by hybridization with an internal neomycin probe produced an 11.1 kb band from the targeted allele and allowed detection of random integration events. Digestion with EcoNI and hybridization with a 519 bp 3’ external probe produced 9.5, 8.4 and 6.8 kb bands from wild-type, targeted and floxed alleles, respectively. Following digestion of genomic DNA samples with the desired restriction enzyme, separation on agarose gel and transfer to nylon membranes, blots were prehybridized and hybridized in 4X SSC, 1 % SDS, 0.5 % skim milk, 20 mM EDTA, 100 μg/ml herring sperm DNA at 65°C for 18 h. Membranes were washed twice in 3X SSC, 1% SDS at 65°C for 15 min, and twice in 0.5X SSC, 1% SDS at 65°C for 15 min. Membranes were exposed for 3 days to BioMax MS film with BioMax intensifying screens.

### Generation of the new floxed *Poldip2* line

The B11-G2 clone of ES cells provided by genOway was tested using a copy number variation assay and appeared to be free from the unwanted duplication that affected clone B12-H5. Thus, clone B11-G2 was used by the Emory Mouse Transgenic and Gene Targeting Core for injection into blastocysts and implantation. Six males with chimerism greater than 70% were produced and were crossed with C57BL/6J females. Two male chimeras were also crossed with Flp deleter females (The Jackson Laboratory, stock number 012930) to delete the neomycin cassette and produce the desired floxed *Poldip2* mice.

### Generation of the excised *Poldip2* line (*Poldip2* knockout line)

Females from the new floxed *Poldip2* line were crossed with a male transgenic carrying EIIa Cre, which is expressed in early embryos (The Jackson Laboratory, stock number 003724). The resulting mosaic pups carried three alleles: wild type, floxed and excised *Poldip2*. Backcrossing to C57BL/6J produced the desired excised (knockout) *Poldip2* heterozygotes.

The three new mouse lines described here have been added to the Mouse Genome Informatics database http://www.informatics.jax.org/:

1. Original floxed *Poldip2* line, with duplication: Dp(11Pigs-Nlk)1Kkgg, MGI:6448073.
2. New floxed *Poldip2* line, without duplication: *Poldip2^tm1.1Kkgg^*, MGI:6448070.
3. Constitutive *Poldip2* knockout line: *Poldip2^tm1.2Kkgg^*, MGI:6448071.

All animal experiments were approved by the Emory University Institutional Animal Care and Use Committee.

### Preparation of genomic DNA

Crude lysates for conventional PCR genotyping were prepared from ear biopsies or yolk sacs using DirectPCR Lysis Reagent (Viagen Biotech). Genomic DNA was purified using the Xtreme DNA isolation kit (Isohelix) or DNeasy Blood & Tissue Kit (Qiagen). These kits were used to purify DNA from crude lysates or from tissue samples preserved in RNA*later* solution (Sigma).

### Preparation of RNA and cDNA

Tissue samples stabilized in RNA*later* solution (Sigma) were homogenized using a TissueLyser II instrument (Qiagen), in the presence of Reagent DX antifoam solution and RLT Plus buffer (Qiagen).

Total RNA was purified using the RNeasy Plus kit (Qiagen). Reverse transcription was carried out using ProtoScript II enzyme (New England Biolabs) and random 15-mer primers. The resulting cDNA samples were purified using the QIAquick PCR purification kit (Qiagen).

### PCR for genotyping and cloning

Primers designed using Oligo software version 7 (Molecular Biology Insights) were ordered from Genosys/Sigma. Genotyping was carried out by conventional PCR using crude DNA lysates and Taq polymerase (New England Biolabs). PCR products destined for cloning were made with Q5 Hot Start High-Fidelity DNA Polymerase (New England Biolabs).

### DNA cloning

Long PCR products were purified by gel electrophoresis and short ones with the QIAquick PCR purification kit (Qiagen). Terminal adenine overhangs were added using Taq polymerase in the presence of dATP, immediately before TOPO T/A cloning. PCR products were cloned into the pCR-XL vector using the TOPO XL PCR cloning kit (Invitrogen). Chemically competent Top10 E. Coli (Invitrogen) were used for transformation.

### DNA sequencing

Standard fluorescent Sanger sequencing was performed by Agencourt/Beckman Coulter or Macrogen, using custom or vector primers and plasmid DNA purified with the QIAprep Spin kit (Qiagen). Reads were assembled using MacVector software.

### Copy Number Variation assays by qPCR

Crude genomic DNA lysates used for routine genotyping were purified before measuring copy number variations by qPCR. Predesigned assays amplifying *Poldip2* at exons 4 or 10, or *Tmem199* at exon 6, as well as *Transferrin receptor 1* (reference gene with a single copy per haploid mouse genome) at exon 17, were purchased from Applied Biosystems. Reactions were performed using TaqMan Genotyping Master Mix, in a StepOne Plus real-time thermocycler (Applied Biosystems). The gene of interest and the reference gene were detected simultaneously in each sample, using TaqMan probes labeled with FAM and VIC, respectively. Quantification was performed using ΔΔCt and CopyCaller software (Applied Biosystems). Copy numbers of the gene of interest are expressed relative to the reference gene and normalized to an average of 2 copies per genome in wild type (diploid) samples.

### qPCR for gene duplication mapping

To facilitate the design of specific genomic primers for duplication mapping, genomic sequence retrieved from the NCBI database was first processed with RepeatMasker software (http://www.repeatmasker.org/). Following primer design with Oligo software, specificity was further evaluated using BLAST software (NCBI). For each primer pair, annealing temperature was optimized experimentally, and quantitative performance verified using serial dilutions of genomic DNA. After optimization, quadruplicate reactions were carried out in a LightCycler instrument (Roche) with glass capillaries, using 5 ng purified genomic DNA, 4 mM MgCl^2^, 300 nM primers, Platinum Taq DNA polymerase (Invitrogen) and SYBR green. A difference in Ct of about 2 cycles between known wild type and homozygous samples was observed for primer pairs annealing near *Poldip2* on chromosome 11. In contrast, there was no difference in Ct for primer pairs annealing far from *Poldip2* or within the leptin receptor on chromosome 4. Thus, the duplication was mapped using numerous custom primer pairs.

### Western blotting

Samples of liver tissue preserved in RNAlater solution were transferred to 500 μl lysis buffer (0.3 M NaCl, 0.2% SDS, 0.1 M Tris base, 1% Triton X-100, 10 μg/ml aprotinin, 10 μg/ml leupeptin, 1 mM PMSF, Halt phosphatase inhibitor cocktail (Thermofisher). Following homogenization (Qiagen TissueLyser II), sonication and centrifugation (20,000 x g) at 4°C for 30 min, pellets were discarded. Equal quantities (20 μg) of protein (Bio-Rad Bradford assay) were brought to equal volumes with Laemmli buffer before boiling at 100°C for 10 min. Proteins were separated using SDS-PAGE and transferred to 0.45-μm pore size PVDF membranes (Millipore). Membranes were blocked in 5% BSA for 1 h before overnight incubation with primary antibodies diluted in Tris-buffered saline with 0.1% tween (TBST). Primary antibodies were: anti-POLDIP2 rabbit monoclonal (Abcam; Ab181841; 1:2000), and anti-tubulin rabbit polyclonal (Abcam; Cat No. Ab6046; 1:5000). Membranes were rinsed with TBST for 30 min before incubation with horseradish peroxidase-conjugated anti-rabbit secondary antibody (Cell Signaling; 7074S, 1:5000) for 1 h. Bands were visualized using enhanced chemiluminescence (ThermoFisher; 34580) and detected with Hyperfilm (Amersham GE).

### RT-qPCR assays of gene expression

Samples of cDNA prepared from mouse tissues were analyzed either with a LightCycler instrument (Roche) with glass capillaries, using Platinum Taq DNA polymerase and SYBR green, or with a QuantStudio 7 instrument (Applied Biosystems), using Forget-Me-Not EvaGreen qPCR Master Mix with low ROX (Biotium). Data quantification was performed using the mak3i module of the qpcR software library (version 1.4-0) [43, 44] in the R environment [45]. Several housekeeping genes were used to normalize the results (β-2 microglobulin, hypoxanthine phosphoribosyl transferase 1 and peptidylprolyl isomerase A), as described earlier [12]. Primer sequences can be found in Table 1.

### RNA Sequencing

#### Tissue preparation ond isolation of RNA

Following euthanasia, mouse carotid arteries were excised after perfusion with normal saline solution and transferred to a 35-mm dish containing ice-cold HBSS. Endothelial-enriched RNA was quickly flushed using an insulin syringe fitted with a 29-gauge needle filled with 150 μL Qiazol solution [46]. Total RNA was purified using the miREasy kit (Qiagen) from both endothelial and leftover (medial) fractions. RNA integrity was assessed with the RNA Nano 6000 assay kit and the Bio analyzer 2100 system (Agilent).

#### Library construction and sequencing

RNA processing was performed using a low-input RNA-seq pipeline from Novogene. Libraries were generated with the NEBNext Ultra RNA Library Prep Kit for Illumina (New England Biolabs). Briefly, mRNA was enriched from 50 ng total RNA using magnetic poly-T beads. First and second strand cDNA was synthesized using random hexamer primers, M-MuLV reverse transcriptase, DNA polymerase I and RNase H, followed by conversion of overhangs to blunt ends. DNA fragments were ligated with NEBNext adaptors and size-fractionated with the AMPure XP system (Beckman Coulter) before treatment with USER enzyme (New England Biolabs) and PCR amplification with universal and index primers using Phusion high-fidelity DNA polymerase. PCR products were purified with the AMPure XP system, and the quality of the library was assessed using the Bio analyzer 2100 system (Agilent). Pooled libraries were read using an Illumina NextSeq 550 sequencer.

#### Read mapping and quantification of gene expression level

Raw reads were processed using in-house Perl scripts at Novogene to remove low-quality data. Clean reads were used to calculate the Q20, Q30 and GC contents. Reads were mapped to genes in the reference mouse genome (mm9) and assembled into transcripts whose abundance was estimated as expected number of Fragments Per Kilobase per Millions base pairs sequenced (FPKM) [47]. Bowtie (v2.2.3) was used to build an index of the reference genome, and TopHat (v2.0.12) to align paired-end clean reads.

#### Differential gene expression analysis

The DESeq2 R package (v1.18.0) was used to compare pairs of sample groups that included four biological replicates. DESeq2 uses a model based on the negative binomial distribution. P values were adjusted with the method of Benjamini and Hochberg to control the false discovery rate [48]. The threshold for significant differential expression was set to q < 0.05.

#### Gene Ontology analysis

Differentially expressed genes were further analyzed using the GOSeq R package. A hypergeometric test, capable of adjusting for gene length bias, served to map genes to terms in the GO database. The false discovery rate was set to less than 0.05. The resulting lists were curated by hand to exclude GO terms unrelated to vascular biology. Finally, GO terms were regrouped into broader categories using the REVIGO software [49].

The RNA-seq results generated in the present study have been deposited in NCBI’s Gene Expression Omnibus [50] database and are available through GEO Series accession number GSE165274. https://www.ncbi.nlm.nih.gov/geo/query/acc.cgi?acc=GSE165274.

#### Statistics

Calculations were performed using GraphPad Prism software versions 7 or 8. Comparisons of observed and expected genotype distributions were performed using two-tailed χ^2^. Comparison of RNA and protein expression between genotypes was performed using two-way ANOVA, followed by Dunnett tests.

## Results

A classical gene targeting strategy in embryonic stem (ES) cells was designed by genOway (Figure 1A) to create a conditional *Poldip2* knockout mouse model. Although it is often recommended to target the first coding exon of a gene, which is exon 1 in *Poldip2*, it seemed safer to flox exon 4 to avoid affecting *Tmem199* expression. Indeed, the *Tmem199* gene is located immediately upstream of *Poldip2* on chromosome 11 (Chr11) and in reverse orientation. Thus, their first exons are only separated by 128 nucleotides and the promoter of *Tmem199* extends inside the *Poldip2* gene. In contrast, the fourth exon of *Poldip2* is separated from the first exon of *Tmem199* by 4,651 nucleotides. Importantly, exon 4 is suitable for targeting because deletion of its 97 bp of coding region would produce a frame shift in the mRNA and a premature stop codon in exon 5.

**Figure 1.**
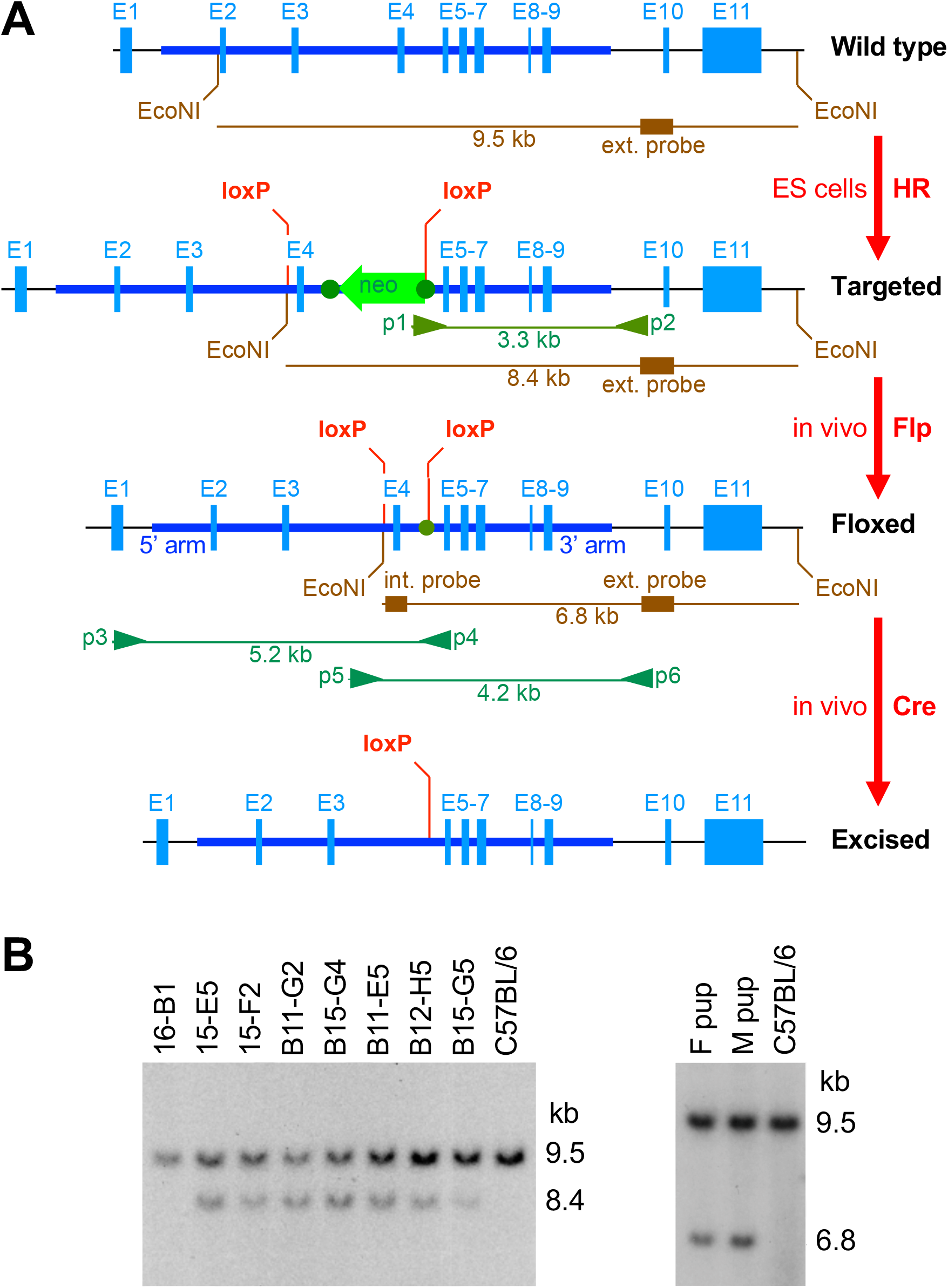
*Poldip2* gene targeting strategy. **A:** Successive modifications of the mouse *Poldip2* gene are shown from top to bottom. The wild type allele from the NCBI database (GRCm38.p6) includes 11 exons, represented by solid blue boxes E1-11. A classical homologous recombination (HR) strategy with a suitable targeting construct was used to flank exon 4 with loxP sites in embryonic stem (ES) cells. Following recombination, the targeted allele includes a neomycin cassette (neo) in antisense orientation in intron 4, surrounded by FRT sites (dark green dots), as well as loxP sites located in introns 3 and 4 (outside neo/FRT). Targeted ES cells were used to produce live chimeric mice. The floxed allele results from in vivo recombination of FRT sites and deletion of the neomycin cassette, after crossing with a Flp deleter transgenic line, thereby generating heterozygous floxed mice. Lastly, the excised allele, which is expected to inactivate *Poldip2*, is produced by recombination of loxP sites and deletion of exon 4, after crossing with Cre transgenic mice. In these maps, the region of homology between the wild type gene and the targeting construct is represented by a thick horizontal dark blue line, extending from intron 1 (5’ arm of the construct) to intron 9 (3’ arm of the construct). DNA segments used as internal and 3′ external probes in Southern blots are represented by brown boxes, together with expected DNA fragments after digestion with EcoNI endonuclease. The annealing sites of primers p1-p6 and corresponding long PCR amplicons are shown in green. **B:** Verification of recombination by Southern blotting. Genomic DNA samples were digested with EcoNI, before electrophoretic separation, blotting and hybridization with the 3′ external probe. **Left blot:** the 7 clones of ES cells in the middle present bands corresponding to both wild type and targeted alleles, suggesting successful recombination. Clone 16-B1 on the left side and the C57BL/6 control on the right side only carry wild type alleles. **Right blot:** after crossing targeted chimeras with Flp deleter transgenics, heterozygous floxed pups were identified. A female (F) and a male (M) pup carried the expected wild type and floxed alleles. These mice, derived from ES cell clone B12-H5, were used as initial founders.

Following transfection of C57BL/6 (B6) ES cells with the targeting construct, 257 G418-resistant clones were isolated and screened using long PCR spanning the 3′ homology arm with primers p1 and p2 (Figure 1A). Ten positive clones were examined using two rounds of Southern blotting, after digestion with EcoRV and EcoNI, respectively. The first blot was hybridized with a neomycin cassette probe and the second one with a 3′ external probe (Figure 1A), hybridizing downstream of the region homologous to the targeting construct. Three clones, which included random integrations, were rejected in the first round. The last 7 clones seemed to be successfully targeted in the second round (Figure 1B left). Six of these clones (all except B15-G5) were used for blastocyst injection and implantation into pseudo-pregnant females. High-percentage male chimeras were crossed with Flp deleter females to produce heterozygous floxed *Poldip2* pups. The genotypes of two mice, derived from clone B12-H5, were confirmed by Southern blotting with the 3′ external probe (Figure 1B right) and were used as founders.

Two females of the following generation were crossed with B6 males in our laboratory to start our first floxed *Poldip2* colony. Genomic DNA from the first pups was used to optimize two genotyping methods, using traditional PCR with primers surrounding either the 5′ or the 3′ loxP site (Figure 2) and their results were in complete agreement. All crosses between heterozygotes (Het) and wild types (WT) produced pups with genotypes in expected proportions (50/50 WT/Het). In contrast, as shown in Table 2, crosses of Het pairs unexpectedly failed to produce homozygotes (Hom). At the time our results were even more confusing than Table 2 would suggest, because we trusted the genotyping methods and included breeders that seemed to be Het, but could have been Hom (both their parents were Het).

**Figure 2.**
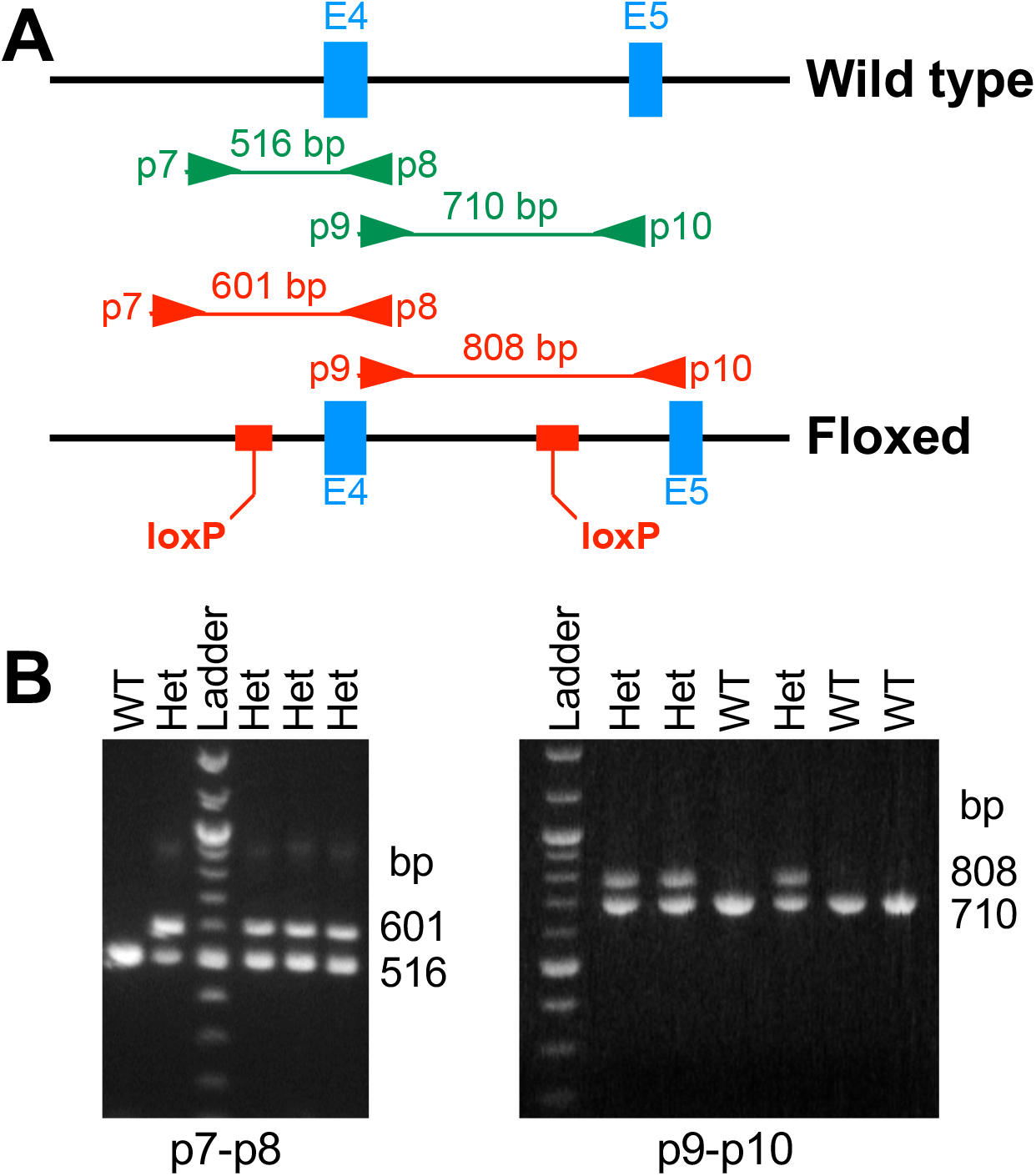
Two floxed *Poldip2* genotyping methods. **A:** Genomic maps of mouse *Poldip2*, around exons 4 (E4) and 5 (E5), showing the annealing sites of PCR primer pairs p7-p8 and p9-p10, surrounding 5′ and 3′ loxP sites, respectively, as well as corresponding amplicons from wild type (green, top) and floxed (red, bottom) alleles. The positions of DNA inserts that include loxP sites are shown in the floxed allele (bottom, red boxes). **B:** Representative agarose gels, following PCR of mouse genomic DNA samples, using primer pairs p7-p8 (left) or p9-p10 (right). Genotypes of individual mice are indicated at the top: wild type (WT) or heterozygous (Het). Steps in the DNA ladder are multiples of 100 bp, with higher intensities at 500 and 1,000 bp.

**Table 2.** **Het x Het crosses of initial floxed *Poldip2* mice fail to produce homozygotes**

| genotype | Observed counts |  | Observed total | Expected total |  |
| --- | --- | --- | --- | --- | --- |
|  | male | female |  | 1:2:1 | 1:3 |
| WT | 13 | 11 | 24 | 30.75 | 30.75 |
| Het | 49 | 50 | 99 | 61.50 | 92.25 |
| Hom | 0 | 0 | 0 | 30.75 |  |
| Total | 62 | 61 | 123 | 123 | 123 |

Although no alteration of function is expected in floxed mice, our previous experience with perinatal lethality in *Poldip2* gene trap mice prompted us to genotype embryos at mid gestation. Thus, we identified two apparently Het male breeders (numbers 82 and 83) which produced 100% (29/29) Het embryos (and no resorption) when crossed with four Het females. Suspecting that these males might really be Hom, we crossed male 83 with 3 WT females, which also produced 100% (21/21) Het embryos. Therefore, this result suggested that male 83 was really a floxed Hom, but also carried WT alleles, which made it appear Het by genotyping.

This interpretation is supported by Table 2, which now only considers litters produced by true Het breeders. We know these breeders had to be Het, not just because of their apparent genotype, but also because one of their own parents was WT. As shown in the rightmost column of Table 2, the distribution of pup genotypes was not different from 1:3, as if Het and Hom genotypes were lumped together.

One possible explanation for this unexpected observation is that either a floxed or a wild type allele was present at a locus different from *Poldip2* in the genome. In addition, because no ectopic integration had been detected in earlier Southern blots, we would expect the genetic defect to have been acquired in a recent cross. In that case it might have been possible to rescue the line, either by identifying animals that had not been affected by the mutation, or by backcrossing to wild type.

To test this hypothesis, two experiments were carried out to verify the genomic context of the *Poldip2* alleles (data not shown). In a first experiment, Southern blots were performed, using genomic DNA samples from a dozen breeders (2 WT, 3 certain Het, and 7 Het that could have been Hom, including male 83). Internal and external probes (shown in Figure 1A) only produced bands expected for both wild type and floxed alleles, confirming that there was no ectopic integration of the targeting construct and implying that all animals, except WT, were Het. Similarly, in a second experiment, long PCRs, spanning either the 5′ or 3′ homology arm, were carried out, using genomic DNA from male 83. These reactions used primers p3-p4 and p5-p6, respectively (Figure 1A: external primers p3 and p6 anneal outside the segment homologous to the targeting construct). The PCR products were cloned and sequenced. All clones were 100% identical to expected alleles, either WT or floxed, including both loxP sites. Thus, both experiments were in agreement and confirmed previous genotyping results: except for two wild types, all mice, including male 83, appeared to be Het, and their alleles seemed to be in the correct genomic context. Furthermore, no unexpected mutation was uncovered by sequencing. Overall, these results were consistent with the possibility that a DNA segment longer than the targeting construct had somehow been duplicated.

To further evaluate the idea of a duplication event, we performed copy number variation assays by qPCR, using commercially available primer and TaqMan probe combinations and included an internal control recommended by the manufacturer. As shown in Figure 3A, genomic DNA samples were tested using primers pairs annealing either inside (*Poldip2* exon 4) or outside the targeting construct (*Poldip2* exon 10), or even in the adjacent gene (*Tmem199* exon 6). As expected, 2 copies per genome were found in wild type mice. In contrast, apparently Het mice fell into two categories: some animals had 3 and others 4 copies per genome at all tested locations. This result supports the interpretation of a long duplication and, together with data above, imply that the floxed allele is associated with an extraneous wild type one. Thus, mice with 3 copies per genome would be true Het and those with 4 copies would be Hom. Interestingly, male 83 carried 4 copies (Figure 3A), as suspected for a functional Hom.

**Figure 3.**
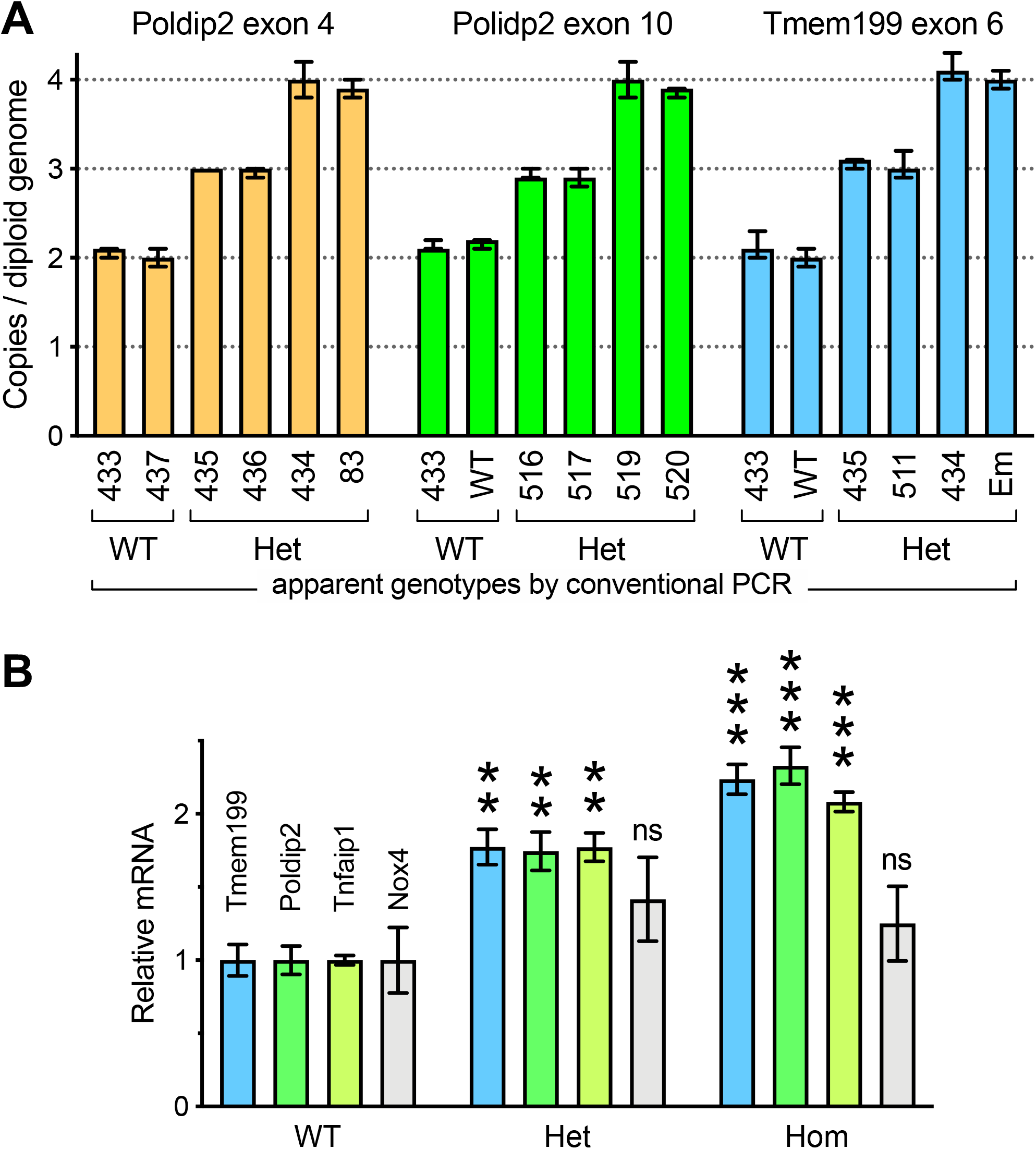
Initial floxed *Poldip2* mice carry a genomic duplication. **A:** Copy number variation assays were performed by qPCR using two primer pairs and two TaqMan probes in each well to amplify both a reference gene and one short genomic segment at the indicated location (top). Bars are averages from quadruplicate representative samples ± confidence intervals calculated by CopyCaller software. Individual mouse ID numbers are indicated below each bar. WT is a wild type control and Em is an embryo. Apparent genotypes, obtained by conventional PCR as in Figure 2, are indicated at the bottom. The presence of 3 and 4 copies per genome is consistent with a duplication carried by heterozygotes (Het) and homozygotes, respectively. **B:** Increased expression of *Poldip2* and neighboring genes in floxed mice. *Poldip2* genotypes were determined by conventional PCR as in Figure 2, in addition to copy number variation assays, as in panel A, assuming that homozygotes (Hom) have 4 copies per genome. Total RNA was purified from duodenum. Expression of *Poldip2* and adjacent genes *Tmem199* and *Tnfaip1*, as well *Nox4*, a distant gene used as a control, was assessed by RT-qPCR and normalized to the wild type for each gene. Bars represent average ± SEM (n= 3-5 mice) **P< 0.01, ***P<0.001 vs. WT.

If a long duplication really occurred and if duplicated genes are functional, one would expect that an increased expression would also be detectable. In tissue samples from mice genotyped using the above copy number variation assays, RNA expression was indeed progressively increased in Het and Hom mice, compared to WT controls (Figure 3B). This change affected not only *Poldip2*, but also both upstream and downstream adjacent genes *Tmem199* and *Tnfaip1*. In contrast, the expression of *Nox4*, a gene located on chromosome 7 was not changed. Thus, so far, the data were consistent with an unexpected long duplication event but did not tell us how far it extended on Chr11 and whether the line could be rescued.

To begin answering these questions, we mapped the duplication by comparing a WT to a Hom sample, using SYBR green qPCR assays. As shown in Figure 4A, primer pairs annealing 28 kb upstream or 52 kb downstream of exon 4 in *Poldip2* readily discriminated between the two samples, indicating that they amplified segments inside the duplication. In contrast, primers annealing within the leptin receptor gene in chromosome 4 amplified both samples at the same cycle number and served as a negative control. Using successive rounds of primer design and testing, we were able to locate the edges of the duplication within 1 kb on each side (Figure 4B). The genomic map in Figure 4C shows that the duplication was much longer than anticipated at 305 kb. It extended further upstream than downstream of *Poldip2* and included at least 15 known genes.

**Figure 4.**
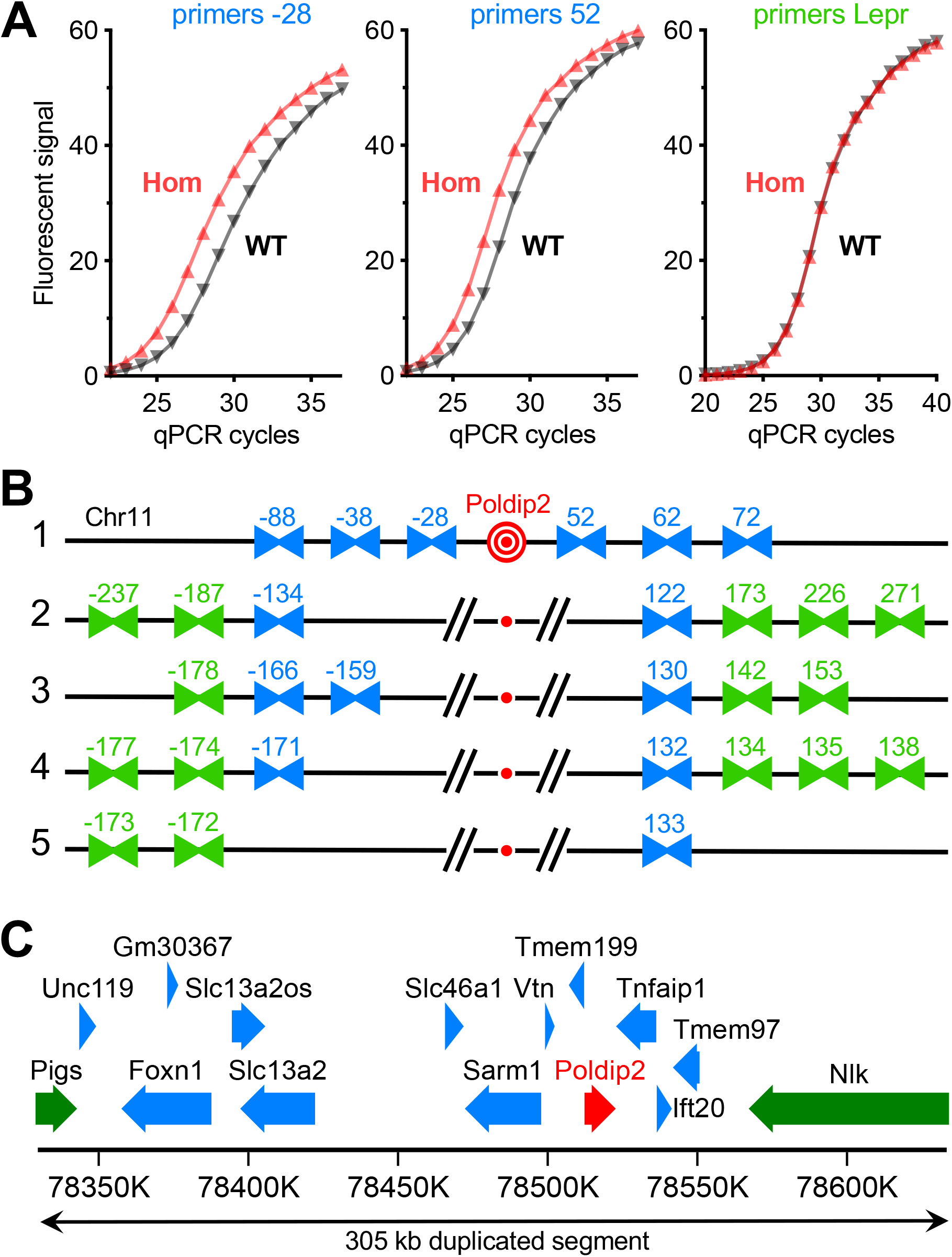
Mapping the genomic duplication in initial floxed *Poldip2* mice. Two samples of genomic DNA were selected to map the duplication: one wild-type (WT) and one homozygous (Hom) floxed *Poldip2*, identified by copy number variation assays as in Figure 3A. Multiple primer pairs, annealing to chromosome 11 (Chr11) on either side of the *Poldip2* gene, were designed. **A:** Validation of the qPCR method with SYBR green detection. The Hom sample amplified earlier than the WT with primer pairs annealing either 28 kb upstream (−28) or 52 kb downstream (52) of exon 4 in *Poldip2*, as expected for duplicated DNA. In contrast, both samples amplified at the same cycle number with a primer pair annealing within the leptin receptor (Lepr) on chromosome 4. Each amplification curve represents the average fluorescent intensity from 4-6 replicates. **B:** Mapping was carried out in five successive rounds of primer design and testing. In this schematic (not to scale), primer pairs are represented by facing arrowheads labeled according to the approximate distance in kb between their annealing site and floxed *Poldip2* (in red at the center). Blue and green colors indicate primer pairs found to anneal inside and outside the duplication, respectively. In the first two rounds of screening, primer pairs were designed to anneal progressively further away from *Poldip2* on Chr11. After finding primers annealing outside the duplication, the last three rounds progressively identified primers closer to the edges of the duplication. In the end, these results indicate that the length of the duplicated segment is between 304 kb (133+171) and 306 kb (134+172). **C:** Genomic map of duplicated DNA segment in floxed *Poldip2* mice. The largest protein coding genes and their orientations are represented as thick arrows. Gene information and nucleotide numbering on Chr11 are from NCBI’s mouse genome (GRCm38.p4). *Poldip2* is shown in red and truncated genes that overlap the edges of the duplication are shown in green.

After mapping the duplication, we investigated its structure. We hypothesized that both copies were present next to each other in the same orientation and connected by a junction sufficiently short to bridge by PCR (Figure 5A). Thus, PCR was attempted using genomic DNA samples from various genotypes with primers annealing near the ends of the duplicated segment and facing outward. The reaction was successful with genomic DNA from Het and Hom mice. Cloning and sequencing of the PCR product provided the exact coordinates of the duplication edges, in addition to revealing that the junction was a simple dinucleotide (Figure 5B). This information allowed us to design a robust PCR genotyping method with two primer pairs to detect both the floxed allele and the junction (as a duplication marker) in a single tube (Figure 5C left). Even mice backcrossed with wild type for several generations were affected and the duplication could be traced back to the earliest animals in our colony (Figure 5C right). Finally, after receiving ES cell clones from genOway, genotyping revealed that the duplication was also present in clone B12-H5, from which our mice were derived (Figure 5C right).

**Figure 5.**
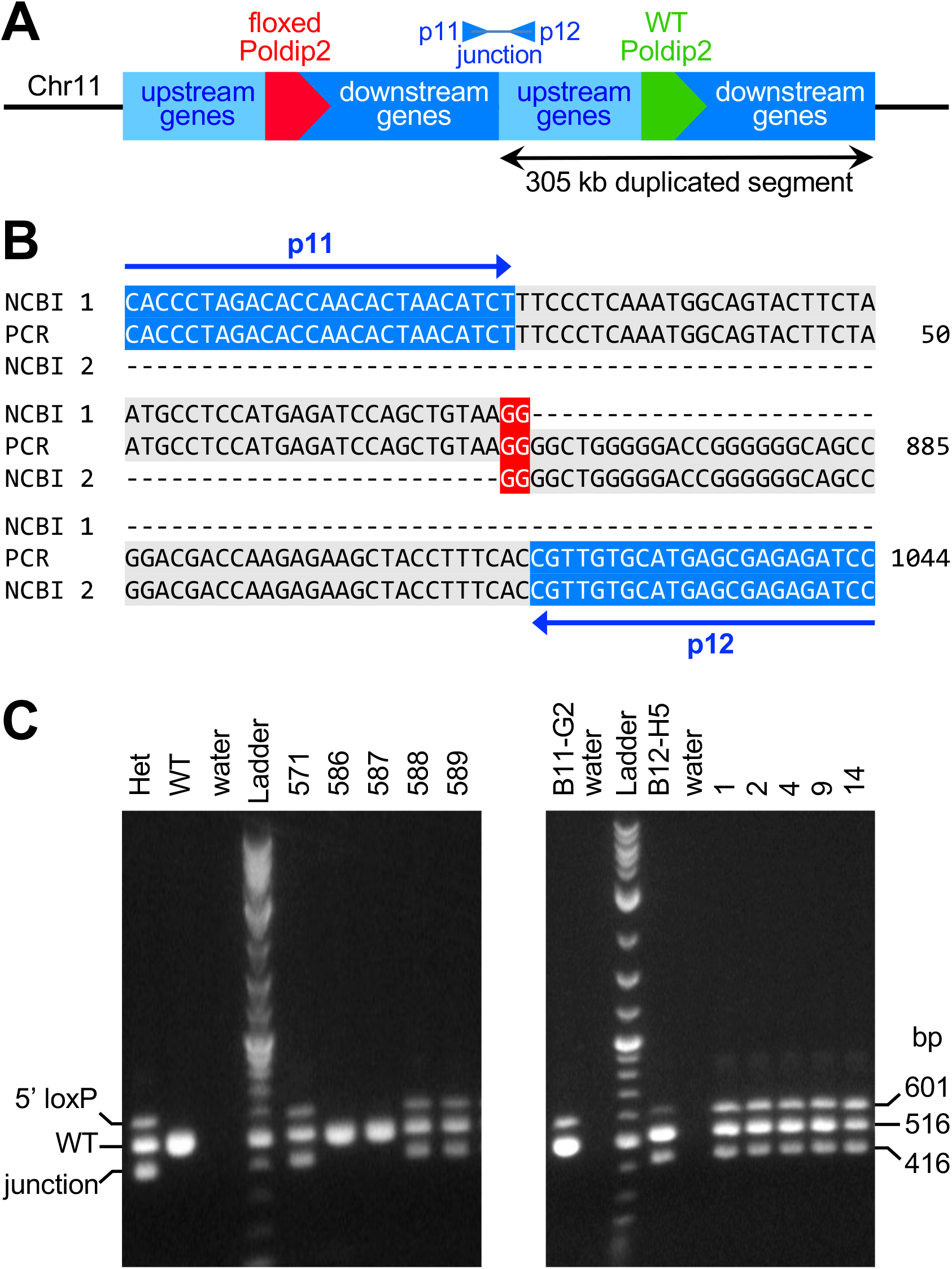
Elucidation of the rearrangement structure provides a new genotyping method. **A:** Schematic of a possible genomic rearrangement with both duplicated segments located side by side in the same orientation and connected by a junction. One copy carries floxed *Poldip2*, while the other retains the WT allele. **B:** Genomic DNA from a homozygous floxed *Poldip2* mouse was amplified by PCR with primers p11 and p12, designed to anneal near the edges of the duplicated segment. A single ~1 kb PCR product was generated, thus confirming the structure proposed in panel A. Following cloning and sequencing of the PCR product, its 1,044 bp sequence was used in a BLAST search of NCBI’s mouse genome. The alignment shows the ends and the center of the PCR product, including primers highlighted in blue and the GG doublet at the junction highlighted in red. Nucleotides 1-862 are 100% identical to a DNA segment (NCBI 1) located downstream of *Poldip2*, while nucleotides 861-1,044 are 100% identical to a DNA segment (NCBI 2) located upstream of *Poldip2*. Sequence coordinates from NCBI GRCm38.p6 (5′ end 78,328,894 and 3′ end 78,634,062) provide the exact length of the duplication: 305,169 bp, thus refining the result from Figure 4B. **C:** A new genotyping method with two primer pairs was designed to detect both the 5′ loxP site (p7 & p8 as in Figure 2) and the duplication junction (p13 & p12) in a single PCR tube. A number of floxed *Poldip2* mice were re-genotyped with this new method with the hope of finding floxed animals without duplication. A representative gel (left) shows that the floxed allele and the mutation were always detected together. Furthermore, the duplication was also found in DNA samples from the earliest Het floxed mice of the colony (right gel, mouse numbers are shown at the top) and in the ES cell clone (B12-H5) from which they derived, indicating that the duplication is stable and occurred during gene targeting. However, another ES cell clone (B11-G2) appeared to be free from the same defect.

In summary, these results show that the undesirable duplication did not appear spontaneously during breeding, but was an accident of gene targeting not readily detectable by Southern blotting, which was transmitted through multiple generations, without resolving itself by recombination with the wild type allele. Therefore, we sought to establish a new line of floxed *Poldip2* mice, starting from ES cell clone B11-G2, which appeared to be successfully targeted and unaffected by the duplication (Figure 5C right and copy number assay not shown). Other ES cell clones shown in Figure 1B were no longer available from genOway and thus could not be genotyped for the duplication.

The Emory Transgenic core facility performed injections of ES cell clone B11-G2 into blastocysts and implantation in pseudo-pregnant females. High-percentage male chimeras were obtained and crossed with Flp deleter females to produce floxed *Poldip2* mice. After verifying by genotyping that the neomycin cassette had been eliminated, we crossed Het mice together. As shown in Table 3, this time Hom floxed *Poldip2* mice were readily identified and the distribution of pup genotypes was not different from 1:2:1, as expected.

**Table 3.**
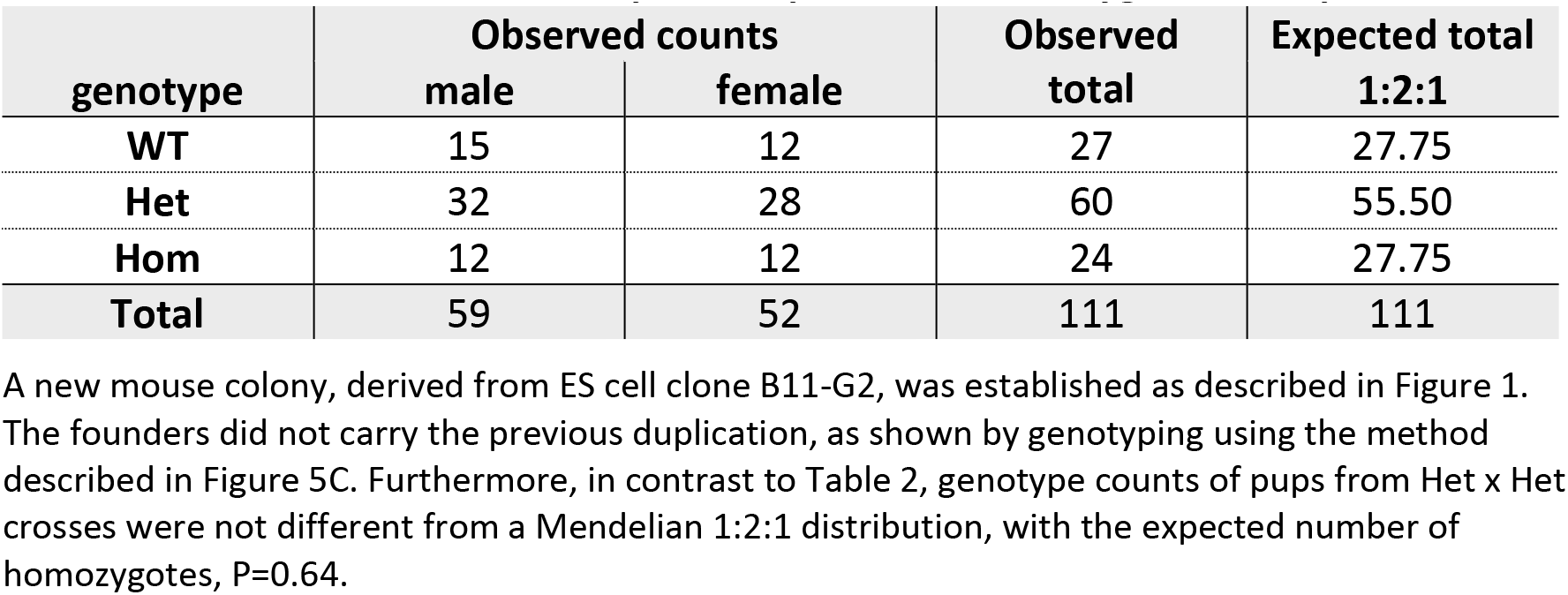
**A new line of floxed *Poldip2* mice produced homozygotes as expected**

Having finally obtained the desired floxed line, it was time to determine experimentally if *Poldip2* is inactivated by Cre-mediated deletion of exon 4. To produce incontrovertible evidence, we first created a new line of mice by crossing floxed *Poldip2* mice with a Cre deleter transgenic. The resulting constitutively excised *Poldip2* Het mice were then backcrossed to B6 WT to remove the Cre transgene from the genome and eliminate mosaicism. To ensure robust genotyping, a new protocol using allelespecific primers was optimized, as shown in Figure 6A. Het mice were then crossed together to determine if it would be possible to produce Hom excised *Poldip2* animals. That was indeed the case, as shown in Figure 6B. Interestingly, Hom mice had no obvious physical deficiency, compared to their wild type siblings.

**Figure 6.**
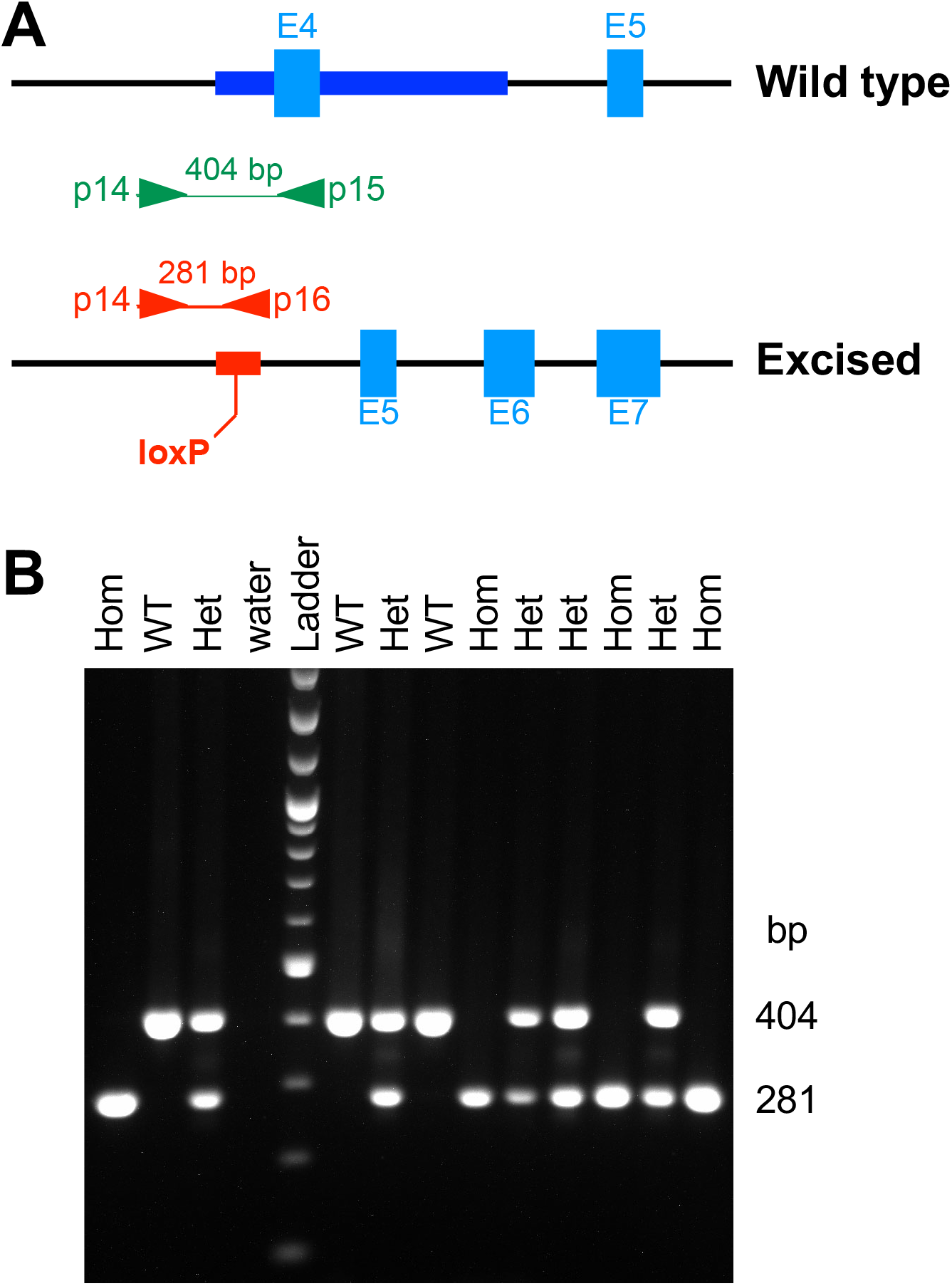
Production of excised *Poldip2* mice. New floxed *Poldip2* mice were crossed with a transgenic Cre deleter to produce animals with a constitutively excised *Poldip2* allele. These mice were then backcrossed with wild type C57BL/6J to remove the Cre transgene and eliminate mosaicism. **A:** Design of a genotyping method for the excised *Poldip2* allele. The genomic maps show the annealing sites of three primers and corresponding amplicons from wild type (top, green) and excised alleles (bottom, red). The 641 bp region around exon 4 (E4) in wild type, highlighted in dark blue, is replaced in the excised allele by a 95 bp insert (red box) that includes the remaining 34 bp loxP site. The forward primer (p14) anneals to intron 3 and the reverse primers either to exon 4 (p15) or to the loxP insert (p16), ensuring allele specificity. **B:** Representative agarose gel, following PCR with the three primers above, revealing the presence of viable homozygous pups at weaning age.

After these necessary preliminaries, we were ready to measure *Poldip2* expression in mice carrying the constitutively excised *Poldip2* allele. As shown in Figure 7A, *Poldip2* mRNA expression in tissue samples was reduced by half in Het and nearly abolished in Hom mice, compared to WT controls, as determined using qPCR. In contrast, expression of the adjacent gene *Tmem199*, was unaffected. It should be noted that excision of exon 4 in *Poldip2* will produce a frame shift and an early stop codon. The resulting message is expected to be unstable and degraded by nonsense-mediated mRNA decay. Therefore, the residual *Poldip2* message detected in Hom mice (Figure 7A) is likely not functional. To further elucidate that point, a Western blot was performed with the same tissue samples. Figure 7B shows that POLDIP2 protein expression was significantly decreased in Het, compared to WT controls, and completely eliminated in Hom mice. Therefore, these results indicate that the excised *Poldip2* allele is null and suitable to produce whole body knockout (KO) mice. These data also give us confidence that crossing floxed *Poldip2* mice with appropriate Cre transgenics will result in the desired cell- or time-specific inactivation of *Poldip2*.

**Figure 7.**
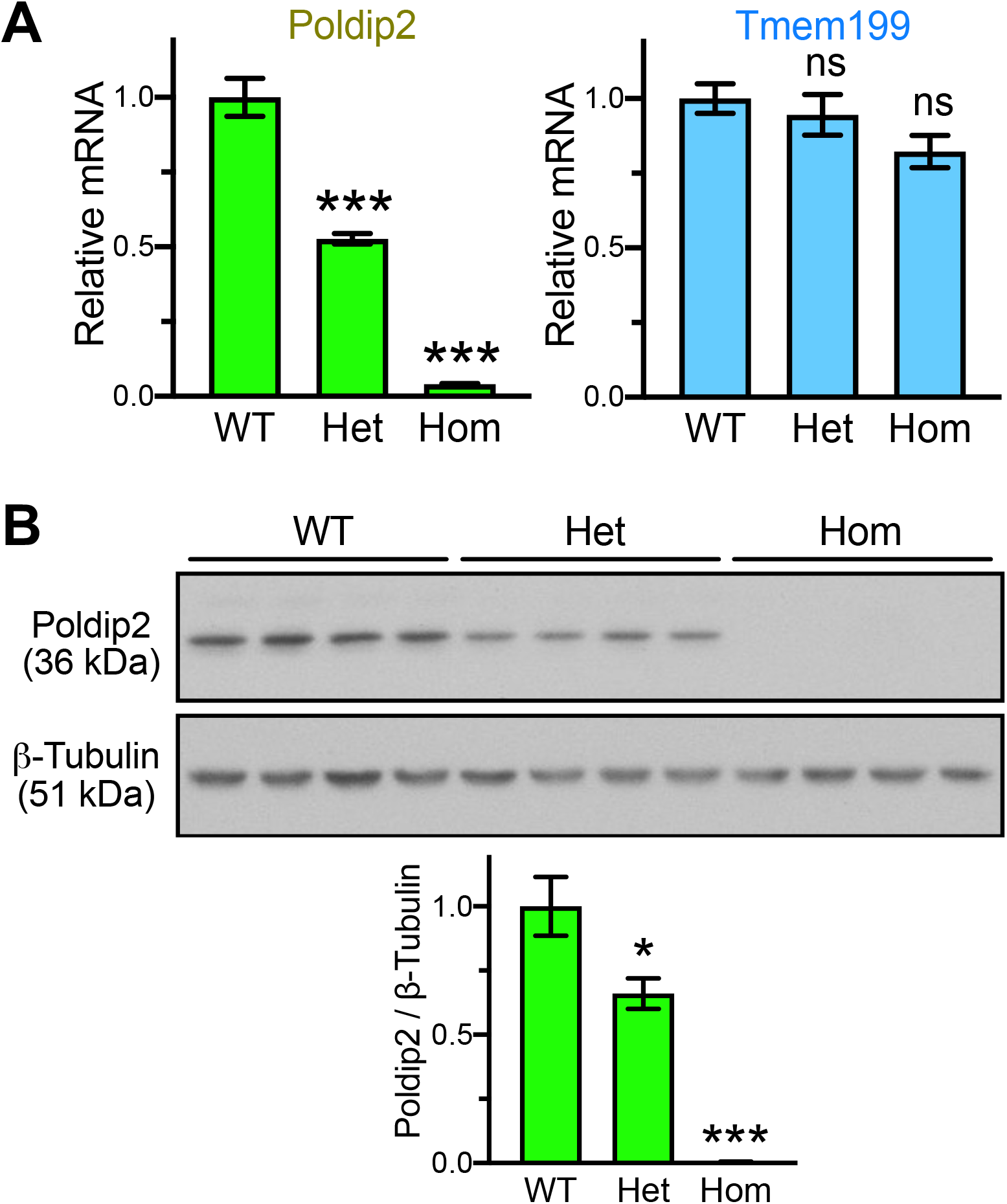
The excised *Poldip2* allele is null and does not affect Tmem199 expression. Gene expression was measured in liver samples from mice with indicated genotypes. Bar graphs represent m ± SEM (n=4 mice in each group). **A:** *Poldip2* (left) and *Tmem199* (right) mRNA expression was measured by qPCR. *** P<0.001 and ns P>0.05, vs. WT. **B:** POLDIP2 and β-TUBULIN protein expression was assessed by Western blotting (top, each lane represents a different mouse) and quantified by densitometry (bottom). * P<0.05 and *** P< 0.001, vs. WT. The absence of POLDIP2 protein expression in Hom mice indicates that they are complete knockouts.

Having shown that it is possible to produce *Poldip2* null mice, we were interested in determining how many live to adulthood, given the lethality observed in *Poldip2* gene trap mice. Table 4 shows that a large proportion of *Poldip2^−/−^* mice were in fact viable. Although the observed distribution of pup genotypes was significantly different from 1:2:1, Hom pups nevertheless represented 18% (55/305) of the total number produced in Het x Het crosses. Thus, the viability of the new line is improved compared to *Poldip2* gene trap mice, which only produced 3% viable Hom pups at weaning age from Het x Het crosses [25].

**Table 4.**
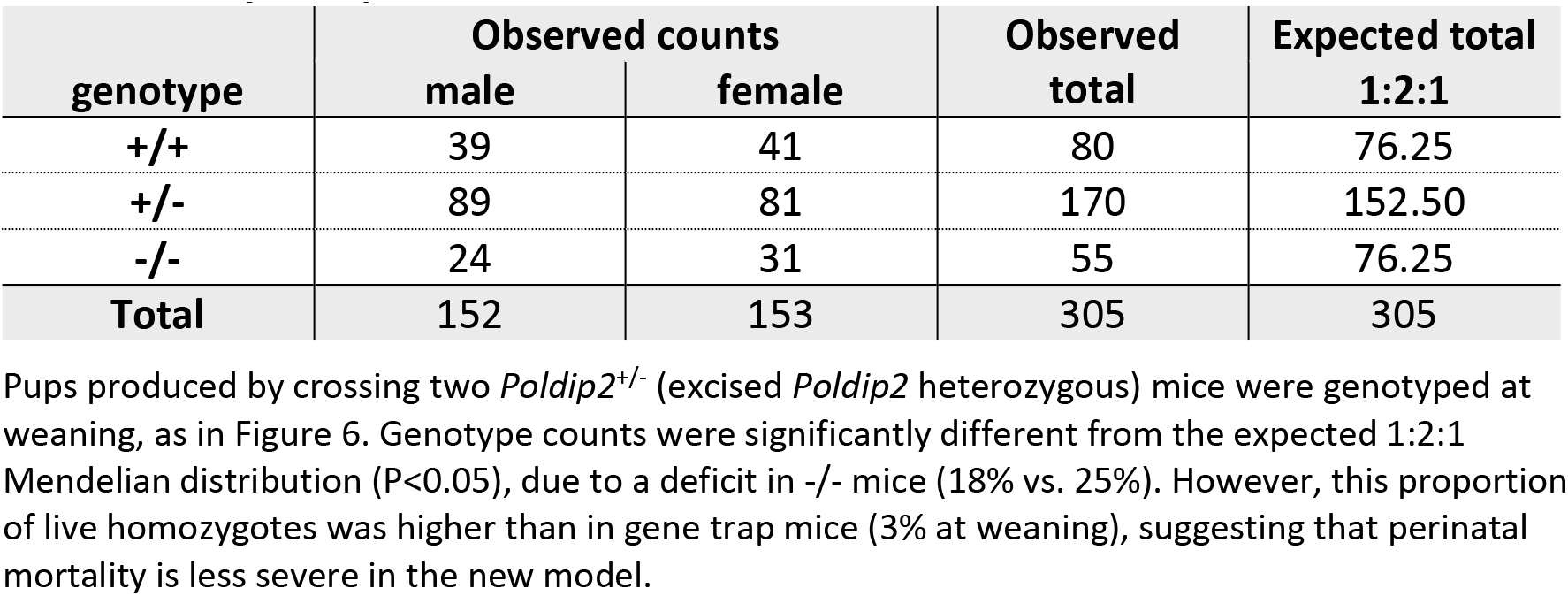
**Many *Poldip2* knockout mice are viable**

To begin assessing the genetic consequences of *Poldip2* deletion in the vasculature, we performed an RNA-seq study. Carotid arteries were isolated from *Poldip2^−/−^* and control mice. RNA was harvested separately from the endothelial layer and the rest of the vessel (labeled medial layer in figures, for simplicity), as described previously [46] and purified from both fractions. Figure 8 shows that the separation of vascular layers was successful, since endothelial and smooth muscle marker genes were almost exclusively expressed in the expected layers, while leukocyte markers were barely detectable, indicating that blood contamination was absent.

**Figure 8.**
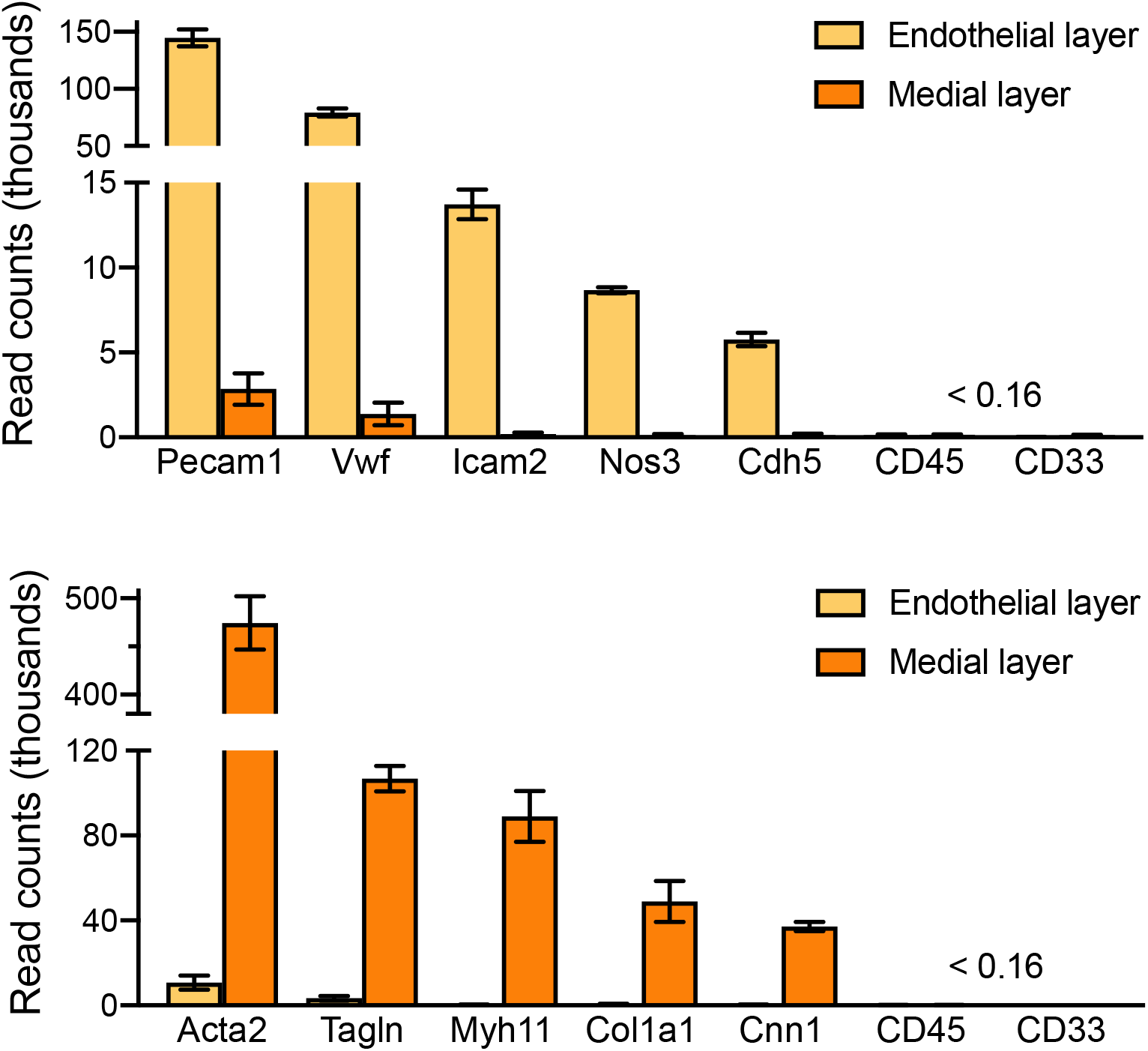
Validation of sample preparation in RNA-seq experiment. RNA was purified from mouse carotid arteries after separation of endothelial and medial layers. The bar graphs represent mRNA expression as average ± SEM of read counts from 4 *Poldip2^−/−^* mice. Similar results were obtained from control mice. Successful separation of vascular layers is indicated by the enrichment in gene markers specific for endothelium (top) and smooth muscle (bottom). Sample contamination by blood was negligible, as shown by the minimal expression of leukocyte markers (CD45 and CD33). Endothelial markers: Pecam1 (platelet endothelial cell adhesion molecule 1), Vwf (Von Willebrand factor), Icam2 (intercellular adhesion molecule 2), Nos3 (nitric oxide synthase 3), Cdh5 (cadherin 5). Smooth muscle markers: Acta2 (smooth muscle alpha actin 2), Tagln (transgelin), Myh11 (smooth muscle myosin heavy chain), Col1a1 (collagen type I alpha 1), Cnn1 (calponin 1).

Next, we examined differential gene expression by RNA-seq in endothelial and medial layers. As expected, a marked decrease in Poldip2 mRNA was observed in knockout mice, compared to controls, in both tissue layers. This is consistent with Figure 7A and supports the notion that deletion of exon 4 produces a non-functional and unstable message. Furthermore, *Poldip2* ablation significantly affected the expression of a large number of genes compared to controls, thus suggesting that it is involved in the direct or indirect regulation of multiple pathways (Figure 9). To investigate this point further, the clusterProfiler software was used to classify differentially regulated genes in both tissue layers. The resulting lists of Gene Ontology (GO) terms were curated manually to remove pathways unrelated to the vasculature and further condensed into broader categories using REVIGO software [49]. The final treemaps presented in Figure 10 are consistent with the known functions of *Poldip2* affecting cell division and migration, the cytoskeleton, the extracellular matrix, and energy metabolism [16, 51]. These results will guide future analysis of the mechanisms of action of POLDIP2 at the molecular level.

**Figure 9.**
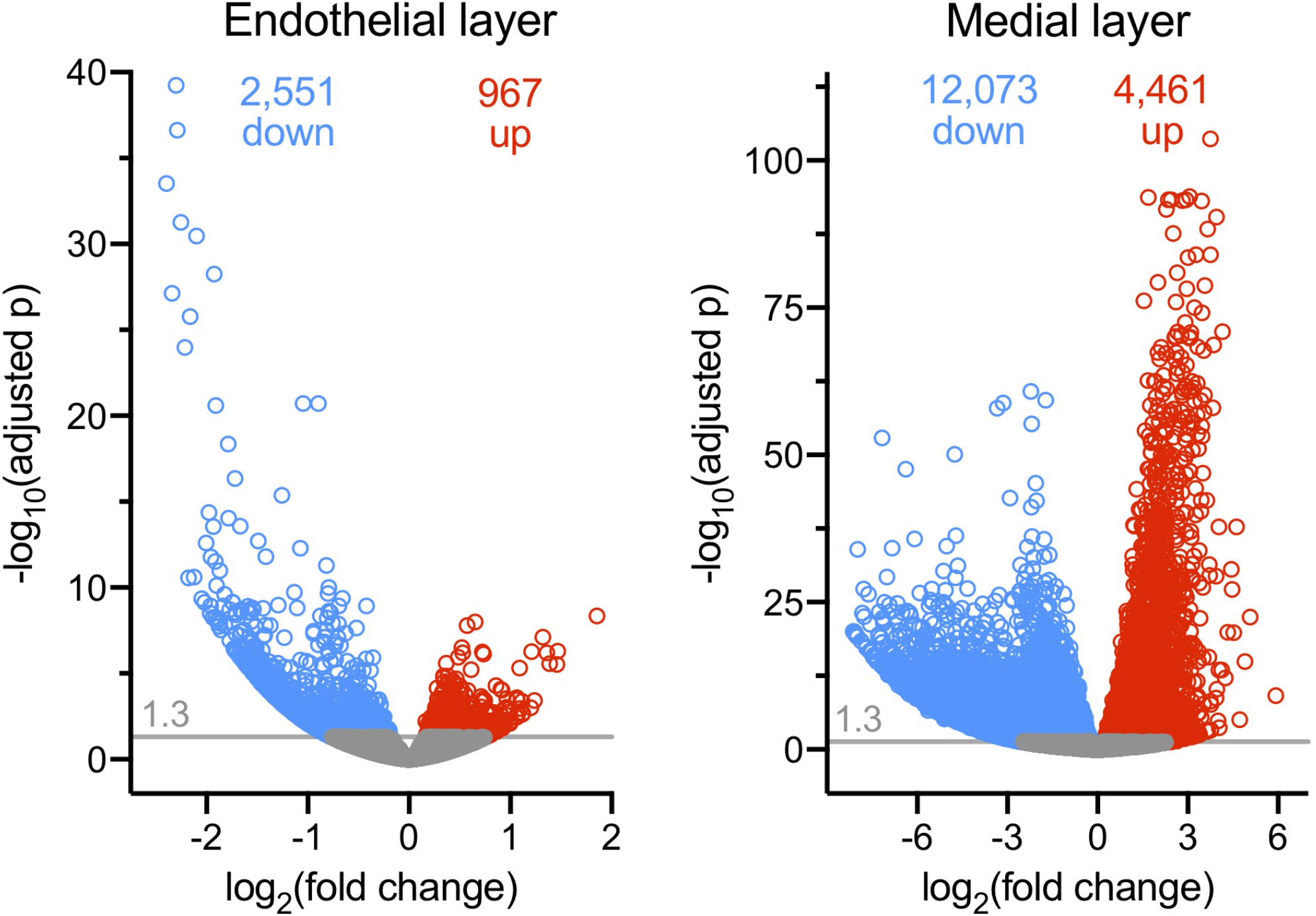
Differential gene expression analysis. RNA-seq data were analyzed with the DESeq2 R package to compare gene expression of *Poldip2^−/−^* and controls (n=4 mice in each group) in the endothelial (left) and medial (right) layers. The threshold for significance of adjusted probability was set at 0.05 (1.3 on the −log10 scale). Genes with significantly reduced or increased expression are indicated by blue and red circles, respectively and their total numbers are indicated near the top of each graph. Genes below the significance threshold appear in gray.

**Figure 10.**
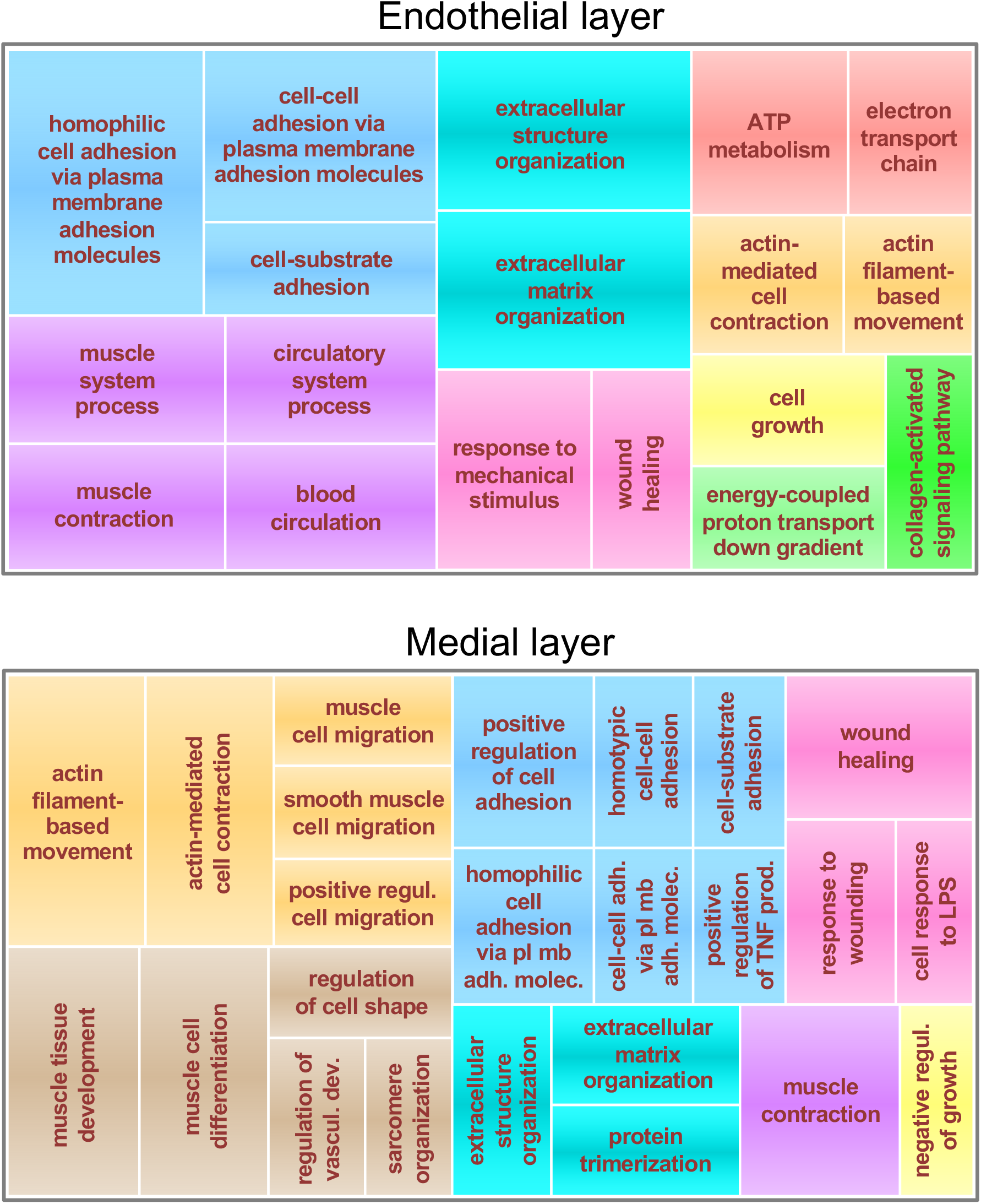
Gene ontology analysis. Genes with significant differential expression in *Poldip2^−/−^* vs. control from endothelial (top) and medial (bottom) layers, were classified using the clusterProfiler software. The resulting lists of GO terms were further summarized using the REVIGO software. The final treemaps represent broad categories of GO terms as boxes with areas proportional to the adjusted probabilities of differentially expressed genes. Related categories are represented in clusters of various colors.

## Discussion

The present study describes the creation of three new lines of mice. First, a line with an unexpected duplication of 15 genes, including different alleles of *Poldip2* (floxed and wild type) in correct genomic contexts. Second, two lines of conditional and constitutive *Poldip2* KO, without duplication. We show that in vivo excision of floxed exon 4 by Cre abolishes *Poldip2* expression, as desired. Furthermore, homozygous constitutive *Poldip2* KO mice could be produced in larger proportions than expected. We have begun characterizing the phenotype of these animals.

Our observation that Het x Het crosses of floxed mice failed to produce Hom pups (Table 2) was baffling because gene expression should not be affected by floxing an internal exon [52]. The problem did not seem to result from incorrect genotyping since two independent methods produced the same results (Figure 2). We began troubleshooting the issue after being assured by the company that there was nothing wrong with the original mice. Thus, we started from the assumptions that a defect must have arisen in an early cross in our laboratory, perhaps due to a genetic instability, and that it should be possible to identify unaffected mice in the colony or rescue others by backcrossing them to wild type.

To address these possibilities, we performed a series of experiments and accumulated evidence pointing to a duplication event that occurred during the earliest stage of the project, in which the floxed allele appeared to be properly recombined, while the endogenous WT allele was retained, making Hom mice appear to be Het by genotyping. In brief: (1) Breeding results from Het x Het crosses (Table 2), suggested that Hom and Het mice were counted together. (2) Male 83 was functionally Hom because it produced 100% Het pups after crossing with WT females. However, three different methods indicated that it was Het: genotyping, Southern blotting and sequencing of long PCR products. (3) In contrast, copy number variations assays (Figure 3A) showed that a number of mice previously thought to be Het, including male 83, were actually Hom, since they carried 4 copies per genome of *Poldip2* and even more surprisingly, also of adjacent gene *Tmem199*. (4) Similarly, mRNA expression (Figure 3B) of *Poldip2* and adjacent genes was progressively increased from WT to Het and Hom. (5) A duplicated 305 kb segment, including at least 15 genes around *Poldip2*, could be mapped by qPCR (Figure 4) with primers annealing at numerous genomic locations. (6) The junction between duplicated segments was amplified and sequenced (Figure 5B), showing that both copies were adjacent and in the same orientation. That information allowed us to design a genotyping method to easily detect the duplication. (7) The duplication was found in the original ES cell clone and in all tested floxed mice (Figure 5C), including those backcrossed to WT for several generations.

The existence of a long duplication explains why it was not detected by long PCRs across homology arms or routine Southern blotting. The purpose of both methods is to verify that the targeted allele is present in the correct genomic context. However, external primers and probes anneal to regions located immediately outside the homology arms (Figure 1A). In our case they produced expected amplicons and Southern band sizes because, although the floxed and the retained wild type allele are present on the same chromosome, they are both surrounded by correct genomic sequence (Figure 5A). Similarly, a classical PCR genotyping method with short amplicons (Figure 2) will detect both alleles, regardless of their chromosomal location.

The observed duplication is much longer than initially suspected. Importantly, it extends well beyond the homology arms of the targeting construct. Both arms cover a 7.3 kb region extending from intron 1 to intron 9 in WT *Poldip2* (Figure 1A), which only represents 2.4% of the length of the duplicated segment (305 kb, Figure 4C). Identifying the cause of such duplications could help prevent similar accidents in future projects. One possibility is that the duplication was already present in ES cells before transfection with the targeting construct. Indeed, cultured ES cell lines can carry karyotype anomalies [53] or smaller duplications caused by replication errors. In that case the targeting construct would have replaced one of the two preexisting WT alleles of *Poldip2*, via a double homologous recombination mechanism. However, this scenario seems unlikely for two reasons. First, the probability of a duplication being present precisely around the gene we were going to target seems low. Second, it would be difficult to explain how that duplication would have affected only part of the initial culture and a subset of the resulting ES cell clones.

A more likely explanation for our observation may be a long extension of the targeting vector, following single-strand invasion of WT sequence. A proposed mechanism, explained in great detail by Mangerich and coworkers [54], as well as the references they cite [55–58], would produce a long double-stranded floxed segment, surrounded by the correct WT genomic context. This long segment would then be inserted randomly elsewhere in the genome. However, because the duplicated and the original segments are present side by side in our case, as well as those of Bismuth et al. [59] and apparently also Mangerich [54], we tend to favor the following hybrid scenario. (1) A single homologous recombination, for example upstream of *Poldip2*, leading to a double-strand break in the targeted allele. (2) Degradation of any vector sequence present at the non-recombined end of the targeting construct. (3) Extension in opposite directions of both construct and targeted allele, along the WT template, thus leading to a long duplication. (4) Non homologous end joining of the newly synthesized segments. It should be noted that the order of the alleles in the duplication represented in Figure 5A is speculative. Our data do not say which of the two alleles is upstream of the other. The floxed allele would end up upstream if the homologous recombination occurred in the 5′ homology arm and vice versa with a 3’ recombination. Elucidation of the exact mechanism of duplication and of the order of alleles in our case would require additional investigations.

Knowing that gene targeting can induce duplications provides an incentive for verifying their absence in targeted ES cell clones before producing live mice. Duplications could be relatively frequent but not often reported, due to the publication bias against negative results. The frequency of duplication events may also be dependent on the locus under investigation. In addition to published examples cited above, in our case we suspect that most of the positive ES cell clones shown in the Southern blot of Figure 1B were similarly affected by duplications. Indeed, in most samples the upper WT band was much more intense than the targeted one. We could not experimentally verify this point because genOway only saved the two clones mentioned in Figure 5C after completion of the project. However, clone B11-G2, with the most balanced band intensity in Figure 1B, was fortunately preserved and turned out to be free from duplication. Another anecdotal piece of evidence regarding the frequent occurrence of long duplications comes from another recent project, carried out for our laboratory by another company, in which a line of knockin mice also failed to produce Hom pups and had to be derived from another ES cell clone. It should be noted that megabase duplications are also observed after repair of double strand breaks induced by CRISPR/Cas9 [60, 61].

We hope our experience will incite others to routinely implement the following recommendations to avoid similar pitfalls. (1) Perform copy number assays of the modified gene in targeted ES cells, or first generation of animals produced using CRISPR/Cas9. The difficulty of reliably discriminating between 2 and 3 copies per genome in Het cells or animals can be alleviated by using several predesigned commercial copy number variation assays near the same locus. Alternatively, digital PCR or comparative genomic hybridization can also detect duplications. (2) Isolate DNA from ES cells alone (without feeder cells) and pay attention to band intensities in Southern blots. Samples in which the WT allele appears to be more abundant than the mutant are suspicious. (3) Simultaneously produce and maintain several independent lines of animals until their genetic modifications have been fully verified. (4) Try to produce homozygotes as soon as possible in a new project. In case of lethality, embryos can be genotyped.

Mice with a large duplication around *Poldip2*, could potentially serve as a useful model for two purposes. First, they could be used to study the effect of increased *Poldip2* expression, compared to control mice in which the extraneous floxed allele would be inactivated with Cre. Second, these mice could also serve as a model of human pathologies associated with amplification of the homologous genomic region on chromosome 17. Multiple records in the ClinVar database are currently associated with pathologies and one publication reports that amplification of *Poldip2* and four adjacent genes correlates with poor prognosis in breast cancer progression [28].

Genotyping for the duplication as in Figure 5C allowed us to show that it was transmitted from the original ES cells to the whole colony and that mice backcrossed to B6 for several generations did not revert to a duplication-free genotype. Because we never found a single floxed mouse in the first colony that did not also carry the duplication and because the B11-G2 ES cell clone appeared to be unaffected, we attempted to use it to produce a new colony, as described in the second part of the present study.

Four lines of evidence indicate that the new floxed *Poldip2* mice, derived from ES cell clone B11-G2, were correctly targeted. (1) The new ES cell clone did not appear to carry the duplication, according to a copy number variation assay (data not shown) and to the new genotyping method (Figure 5C). (2) Similarly, genotyping showed that new floxed *Poldip2* mice did not carry the same duplication as the first line (data not shown). (3) The distribution of genotypes in pups from Het x Het crosses was not different from 1:2:1 (Table 3). Thus, Hom floxed mice were readily identified for the first time. (4) A constitutive KO line, with expected effects on *Poldip2* expression, could be derived from the floxed line (Figure 7).

A large proportion of the new constitutive *Poldip2^−/−^* mice lived until weaning age (Table 4) and beyond. This was unexpected because most Hom *Poldip2* gene trap mice died around birth [12, 25] and *Poldip2* expression is abolished in Hom of both strains (Figure 7 and [25]). This difference in viability between the two strains may be due to decreased *Tmem199* expression in *Poldip2* gene trap mice (data not shown), resulting from a disruption of the *Tmem199* promoter. Indeed, because the two genes are in opposite orientation (Figure 4C) and their first exons are only separated by 13 bp, the gene trap insert in intron 1 of *Poldip2* [12] is also located within the promoter of *Tmem199* (only 901 bp from the start of its first exon). Further investigations of *Tmem199* would be required to assess its effects on viability. Fortunately, *Tmem199* expression is not affected in our new *Poldip2^−/−^* mice (Figure 7A), presumably because the excised exon 4 in *Poldip2* is located 4.6 kb downstream of exon 1, rather than in the *Tmem199* core promoter.

Besides revealing that ablation of *Poldip2* alone is not lethal to the degree observed in the gene trap model, the new knockout line allows experiments to be carried out in Hom animals. Thus, our initial RNA-seq results confirm that POLDIP2 is multifunctional, directly or indirectly, since its deletion affects multiple pathways (Figure 10). Furthermore, the pathways identified here are consistent with previous studies from our laboratory performed in cultured cells [10, 31, 32], which revealed the importance of POLDIP2 in cytoskeletal dynamics. Our results are also consistent with studies carried out in *Poldip2* gene trap mice and cells derived from them [1, 4, 7, 12, 25, 26, 30, 36, 37, 40, 51], which provided insights into the functions of *Poldip2* in the vasculature, in inflammation and mitochondrial function. Therefore, with the exception of homozygous lethality, data provided by the gene trap model are in agreement with the present results. In the future, floxed and *Poldip2^−/−^* mice and genes identified in the present RNA-seq experiment will serve to further elucidate *Poldip2* functions at the molecular level.

In summary, although our project suffered from an initial setback due to an unexpected duplication, our experience can benefit those who want to create knockouts, knockins or generally edit genes in cells or whole organisms. In the end, we were able to produce the desired lines of mice, both conditional and constitutive *Poldip2* knockout. Importantly, we showed that excision of floxed exon 4 with Cre produces a null allele and began characterizing the phenotype of *Poldip2* knockout animals. Both new lines of *Poldip2* mice can be used to further investigate the roles of this multifunctional protein in physiological and pathological conditions.

## Acknowledgements

Southern blots performed at genOway were included with permission from the company.

This research was funded by National Institutes of Health grants HL95070 and HL152167 to K.G., HL119798, HL095070, and HL139757 to H.J., who was also supported by the Wallace H. Coulter Distinguished Faculty Chair Professorship.

This study was supported in part by the Mouse Transgenic and Gene Targeting Core (TMF), which is subsidized by the Emory University School of Medicine and is one of the Emory Integrated Core Facilities. Additional support was provided by the Georgia Clinical & Translational Science Alliance of the National Institutes of Health under Award Number UL1TR002378. The content is solely the responsibility of the authors and does not necessarily reflect the official views of the National Institutes of Health.

